# Evolution of an amniote-specific mechanism for modulating ubiquitin signalling via phosphoregulation of the E2 enzyme UBE2D3

**DOI:** 10.1101/750505

**Authors:** Monica Roman-Trufero, Constance M Ito, Conrado Pedebos, Indiana Magdalou, Yi-Fang Wang, Mohammad M Karimi, Benjamin Moyon, Zoe Webster, Aida di Gregorio, Veronique Azuara, Syma Khalid, Christian Speck, Tristan Rodriguez, Niall Dillon

**Affiliations:** Gene Regulation and Chromatin Group, MRC London Institute of Medical Sciences, Hammersmith Hospital Campus, Du Cane Road, London W12 0NN, UK; Dept. of Chemistry, University of Southampton., Southampton SO17 1BJ; DNA Replication Group, Institute of Clinical Sciences, Imperial College, Hammersmith Hospital Campus, Du Cane Road, London W12 0NN, UK; Bioinformatics and Computing, MRC London Institute of Medical Sciences, Imperial College, Hammersmith Hospital Campus, Du Cane Road, London W12 0NN, UK; Transgenics and ES cell Facility, MRC London Institute of Medical Sciences, Imperial College, Hammersmith Hospital Campus, Du Cane Road, London W12 0NN, UK; BHF Centre for Research Excellence, National Heart and Lung Institute, Imperial College London, Hammersmith Hospital Campus, Du Cane Road, London W12 0NN, UK; Institute of Reproductive and Developmental Biology, Imperial College London, Hammersmith Hospital Campus, Du Cane Road, London W12 0NN, UK

## Abstract

Genetic variation in the enzymes that catalyse post-translational modification of proteins is a potentially important source of phenotypic variation during evolution. Ubiquitination is one such modification that affects turnover of virtually all of the proteins in the cell in addition to roles in signalling and epigenetic regulation. UBE2D3 is a promiscuous E2 enzyme that acts as a ubiquitin donor for E3 ligases that catalyse ubiquitination of developmentally important proteins. We have used protein sequence comparison of UBE2D3 orthologues to identify a position in the C-terminal α-helical region of UBE2D3 that is occupied by a conserved serine in amniotes and by alanine in anamniote vertebrate and invertebrate lineages. Acquisition of the serine (S138) in the common ancestor to modern amniotes created a phosphorylation site for Aurora B. Phosphorylation of S138 disrupts the structure of UBE2D3 and reduces the level of the protein in mouse ES cells (ESCs). Substitution of S138 with the anamniote alanine (S138A) increases the level of UBE2D3 in ESCs as well as being a gain of function early embryonic lethal in mice. When mutant S138A ESCs were differentiated into extra-embryonic primitive endoderm (PrE), levels of the PDGFRα and FGFR1 receptor tyrosine kinases (RTKs) were reduced and PreE differentiation was compromised. Proximity ligation analysis showed increased interaction between UBE2D3 and the E3 ligase CBL and between CBL and the RTKs. Our results identify a sequence change that altered the ubiquitination landscape at the base of the amniote lineage with potential effects on amniote biology and evolution.

## INTRODUCTION

Ubiquitination is an ancient post-translational modification that originated in Archaea (Grau-Bove et al., 2015) and controls a wide range of cellular and developmental processes across eukaryotic lineages (reviewed by (Rape, 2018). One of the major functions of ubiquitination is protein quality control and regulation of the rate of turnover of individual proteins. This occurs through the addition of polyubiquitin chains, which act as signals that direct proteins to the proteasome for degradation. In addition to these functions, ubiquitination is involved in diverse processes that include epigenetic regulation of transcription, control of lysosomal trafficking of growth factor receptors, regulation of DNA repair and modulation of signalling pathways including, among others, the MEK/ERK NF-*κ*B and Wnt signalling pathways. These different types of regulation can occur through a variety of ubiquitination events including monoubiquitination of specific residues and addition of polyubiquitin chains with different topologies generated by varying combinations of ubiquitin chain linkages (Rape, 2018).

Ubiquitin is first activated by formation of a thioester bond with the E1 enzyme, and is then transferred to an E2 ubiquitin conjugating enzyme. The E2 enzyme acts as the donor for transfer to the substrate, a reaction that is catalysed by the action of an E3 ubiquitin ligase, which is the component of the cascade that confers most of the substrate specificity. Mammals have 1 to 2 E1 enzymes, around 30 E2 enzymes and at least 600 E3 ligases. This means that each E2 generally interacts with multiple E3 enzymes. Together with the fact that different E2s preferentially give rise to ubiquitin chains with different linkages (Ye and Rape, 2009), this gives E2 enzymes a broad potential for influencing cellular functions that has still to be explored in depth.

UBE2D3 is a promiscuous E2 enzyme that acts as the donor for a number of E3 ligases that are involved in protein quality control and important signalling and regulatory pathways (Brzovic and Klevit, 2006; Jensen et al., 1995; Wenzel et al., 2011). The UBE2D family members are the vertebrate orthologues of the Drosophila Effete (Eff) protein (also known as UBCD1), which shares 94% identity with human UBE2D3 and has been shown to have multiple roles in *Drosophila* developmen*t* (Chen et al., 2009; Cipressa and Cenci, 2013).

Among the E3 ligases that use UBE2D3 as a ubiquitin donor in vertebrates is the developmentally important CBL proto-oncogene (Liyasova et al., 2019), which is responsible for controlling the endocytosis and lysosomal trafficking of the epidermal growth factor receptor (EGFR) (Fortian et al., 2015) and ubiquitination and degradation of platelet-derived growth factor receptor-α (PDGFRα) and fibroblast growth factor receptors (FGFR) (Haugsten et al., 2008; Vantler et al., 2006). Other E3 ligases for which UBE2D enzymes have been shown to act as ubiquitin donors in vertebrates include the Polycomb protein Ring1B (Bentley et al., 2011), which catalyses mono-ubiquitination of histone H2AK119, MDM2, which ubiquitinates the tumour suppressor and checkpoint protein p53 and regulates its turnover (Saville et al., 2004), and CHIP/STUB1, which plays an essential role in protein turnover and quality control (VanPelt and Page, 2017).

Acquisition of the capacity to live entirely on land was a key event in vertebrate evolution that took place during the Carboniferous period. The transition occurred when tetrapod amphibia that had evolved in the Devonian acquired the ability to breed on dry land without returning to water, leading to evolution of the amniote lineage (reviewed by (Clack, 2012). One of the major changes that led to this capability was the development of extraembryonic membranes, which surrounded the amniote embryo, protecting it from dessication (Ferner and Mess, 2011; Stern and Downs, 2012). The first true amniote fossil has been dated to 314 Mya (Carroll, 1964) and the common ancestor to amniotes is thought to have lived between 340 and 314 Mya (Clack, 2012). Other changes that made a terrestrial existence possible included skeletal alterations that allowed movement and feeding on land (Clack, 2012) and the physiological changes that made it possible for amniotes to become fully air breathing. The detailed molecular mechanisms that led to these changes are still poorly understood but they are likely to have included a mixture of changes to transcription factor mediated control of developmental genes and to the signalling pathways that control development and organogenesis.

Here we describe an unusual mutation that occurred in the common ancestor to modern amniotes generating a single amino acid change at a highly conserved site in the UBE2D3 gene. The substituted serine (S138) is completely invariant across amniote lineages whereas the position is occupied by a conserved alanine in anamniote vertebrates, invertebrates and single-celled eukaryotes. We show that phosphorylation of S138 by Aurora B kinase disrupts the structure of UBE2D3, destabilising it and reducing its level and activity. The reduction in UBE2D3 activity affects the functioning of the CBL E3 ligase, increasing the expression of RTKs in differentiating extra-embryonic primitive endoderm (PrE). Mutation of the S138 residue to the anamniote alanine has a gain of function effect that results in early embryonic lethality in mouse embryos, compromised ability of mutant ESCs to develop into PrE and reduced levels of PDGFRα and FGFR1 in differentiating PrE. Our results identify a novel regulatory pathway that originated in the common ancestor to modern amniotes and affected the ubiquitination landscape with potential impacts on amniote evolution.

## RESULTS

### Comparison of UBE2D3 orthologues across eukaryotic lineages

The amino acid sequence conservation of UBE2D3 orthologues was analysed by carrying out a sequence comparison across 118 eukaryotic species using the human sequence as a reference (Figure 1 and Table S1) (see Methods for details). The results show a very high level of conservation, with 94% sequence identity between humans and *Drosophila melanogaster* and >80% identity observed between humans and basal eukaryotes such as the Oleaginous diatom, *Fistuilifera solaris*, and the extremophilic red alga, *Galdieria sulphuria* (Figure 1 and Table S1). As expected, the overall variation relative to human UBE2D3 increases with increasing evolutionary distance but the variation is non-random and confined to specific residues or short regions while significant stretches of the protein are completely invariant. Many of the substitutions observed are conservative and, even where the substitution is non-conservative the variation is generally restricted to a few amino acids. This suggests that the variable regions have also been subject to significant functional constraint during evolution.

**Figure 1.**
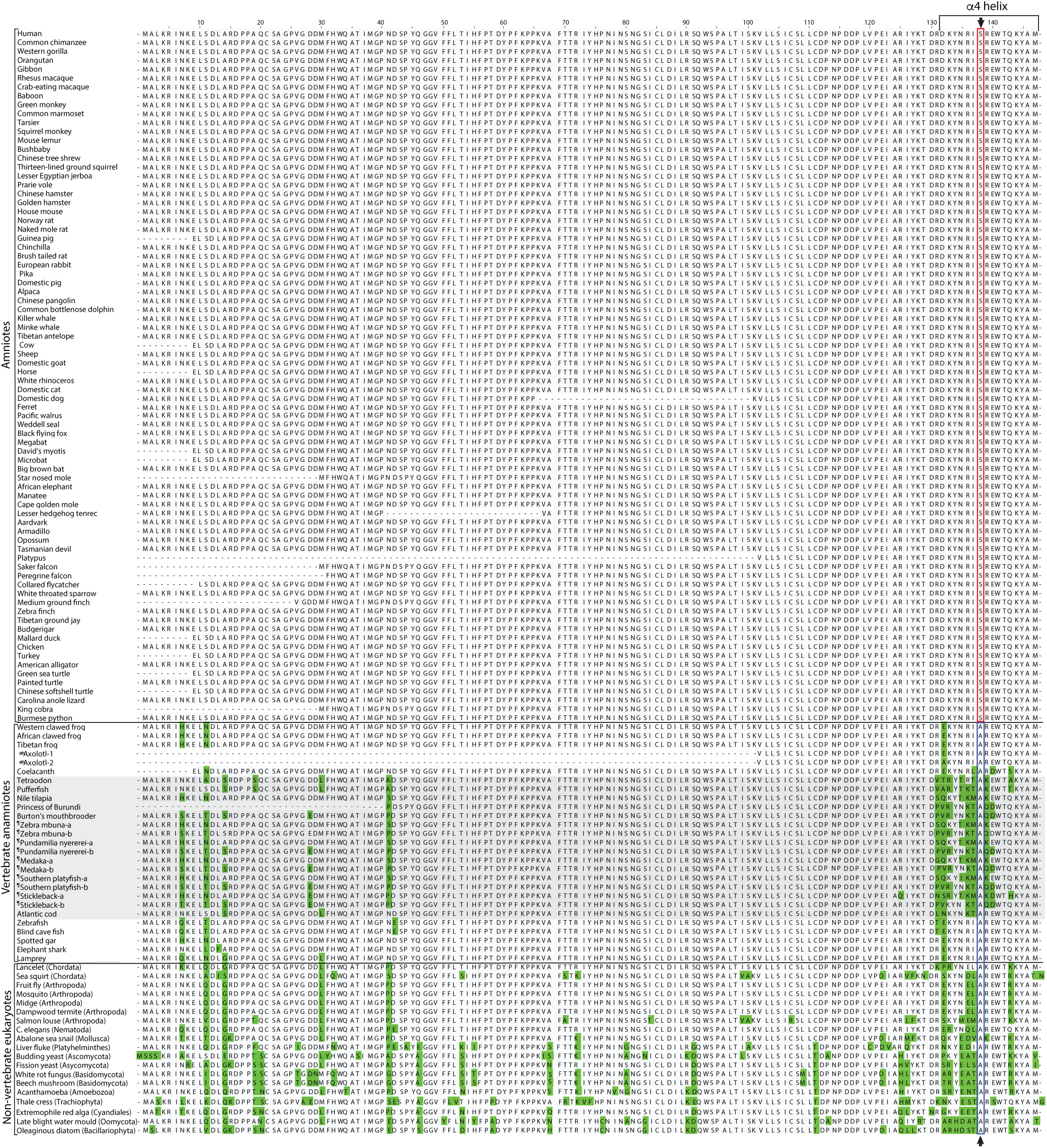
Alignment of the sequences of vertebrate UBE2D3 genes and non-vertebrate UBE2D3 orthologues. Numbering starts from the first residue of the human sequence. Residues that differ from the human sequence are highlighted in green with the exception of position 138. The serine and alanine residues at position138 are indicated by red and yellow boxes respectively. Dashed lines indicate gaps in some of the available sequences. Full species names and accession numbers and sources for the sequences are provided in Tables S1 and S2. #The Axolotl Exon-6 and exon-7 sequences were located for two UBE2D genes by blast searching the Axolotl genome sequence (for details, see Supplementary Methods). Sequence comparison indicated that these genes were UBE2D2 or UBE2D3, but it was not possible to determine which is UBE2D3. Therefore, both sequences are shown as Axolotl-1 and −2. The sequences differ by a single residue (position 132), which is occupied by E or A. ¶Duplicated UBE2D3 genes (UBE2D3-a and UBE2D3-b) arising from the whole genome duplication that occurred in the common ancestor to teleost fish.

The highest variability in the UBE2D3 sequence is observed in a stretch of 20 residues (127-147) that contains an α-helix at the C-terminal end of the protein (Figure 1, Figure S1 and Table S1). The variation within the C-terminal region is also non-random. For example, basal fish lineages (Lamprey, Elephant shark, Spotted gar) show 90% homology between this region and the equivalent region of human UBE2D3, whereas in a subgroup of teleost fish, the Acanthomorpha, the homology ranges from 55-75%, and is lower than the homology between humans and the majority of invertebrate metazoan species examined (Figure 1 and Table S1, grey shading). The Acanthomorpha (spiny rayed fishes) are a successful and morphologically diverse fish taxon that first appeared approximately 120 Mya and comprise around one third of all living vertebrates (Bellwood et al., 2015; Chen et al., 2014; Friedman, 2010). They include the pufferfish, cichlids, sticklebacks and seahorse as well as cod and sea bass. Several of the Acanthomorph species examined had gene duplications that are likely to be a product of the whole genome duplication that occurred in the common ancestor to teleosts (Meyer and Van de Peer, 2005; Opazo et al., 2013). Neo-functionalisation or sub-functionalisation of these duplicated genes could have contributed to the increased variation in the C-terminal region of Acanthomorphs. It is notable that the side-chains of the hypervariable amino acids are mainly located on the outward facing surface of the α4-helix whereas the invariant residues mostly face inwards towards the hydrophobic core of the protein (Figure S1B). This suggests that the variable residues are involved in protein-protein contacts and that these contacts have been subject to evolutionary change in the Acanthamorpha. Additional discussion of the results obtained from the analysis of fish sequences can be found in Supplementary Information.

### Evidence of an alanine to serine substitution at UBE2D3 position 138 in the common ancestor to amniotes

A further striking feature of the C-terminal region of UBE2D3 that was revealed by the sequence comparison was the very high conservation of the alanine residue at position 138 (A138) in anamniotes, invertebrates and single celled eukaryotes (Figure 1, Table S1). The residue occupying position 138 is located in the centre of the C-terminal α4-helix (Figure S1) and the conservation of alanine at this position is maintained even in the Acanthamorph fish, where the residues on both sides of position 138 are highly variable. However, in all amniote species examined, this position is occupied by serine (S138), a residue that is not found at position 138 in any of the other eukaryotic species examined. This leads to the conclusion that an alanine to serine substitution occurred in the common ancestor to modern amniotes and that the S138 residue has been conserved in all amniote lineages. The sequence context around position 138 in amniotes means that the substitution created a consensus phosphorylation site for the Aurora B kinase and we have previously reported that UBE2D3-S138 is phosphorylated *in vitro* and *in vivo* by Aurora B (Frangini et al., 2013).

The variation observed in teleosts in general and the Acanthomorpha in particular, contrasts with the near-absolute conservation of the entire UBE2D3 sequence across Amniote orders (Figure 1). This level of conservation of the amino acid sequence of UBE2D3 has occurred despite the fact that the common ancestors to modern amniotes and teleosts existed at roughly the same time during the Carboniferous period and suggests that the UBE2D3 sequence is subject to a high level of functional constraint in amniotes.

In summary, our comparative analysis of the UBE2D3 sequences leads us to conclude that a unique alanine to serine substitution in the C-terminal region of UBE2D3 occurred in the common ancestor to modern amniotes, creating a consensus phosphorylation site for the Aurora B cell cycle kinase. The occurrence of the substitution at a critical point in vertebrate evolution and its absolute conservation in amniotes raised the possibility that the acquisition of the serine residue at this position played a functional role in amniote evolution.

### Mutation of S138 to the anamniote alanine increases the stability of UBE2D3 in ESCs

Analysis of the mRNA levels of UBE2D family members in pluripotent mouse ESCs showed that UBE2D3 is the predominantly expressed UBE2D enzyme in these cells with UBE2D2 and UBE2D1 showing barely detectable levels of mRNA (Figure S2A). This allowed us to investigate the functional roles of the UBE2D3-S138 residue in the embryonic cell lineages of a model amniote by replacing it with the anamniote alanine residue in mouse ESCs using CRISPR/Cas9. Following transfection of ESCs with the guide RNA (gRNA), Cas9 expression construct and targeting sequence (see Methods), clones were isolated and genotyped by nested PCR. Out of a total of 37 clones analysed, 10 were homozygous for the mutation. The mutant cells grew normally and showed no overt changes in cell cycle profile (Figure S2B). To ensure that any differences in the behaviour of the cells were caused by the mutation, CRISPR/Cas9 mutagenesis was used to generate a revertant cell line from one mutant clone. For subsequent functional analysis of the ESCs, wild-type ESCs were compared with the S138A mutant clone and the A138S revertant that was generated from it.

When the mutant UBE2D3 protein was analysed by western blotting of protein extracts from the UBE2D3-S138A ESCs, it became apparent that the presence of the S138A mutation enhanced the stability of the protein (Figure 2A). The mean level of UBE2D3 was increased by approximately 3-fold (P<0.05, n=5). Analysis of UBE2D3 mRNA levels showed no significant change in the transcript levels in the mutant and revertant cells compared with wild-type cells (Figure S2C). Inhibition of protein synthesis by incubation of wild-type and mutant cells with cycloheximide showed a significantly lower rate of degradation of UBE2D3-S138A compared with wild-type UBE2D3 after 5 hours of cycloheximide treatment, confirming the increased stability of the mutant protein (Figure 2B). Our findings provide evidence that the alanine to serine substitution at position 138 substantially affected the stability of UBE2D3, thereby altering the functioning of the protein in amniotes.

**Figure 2.**
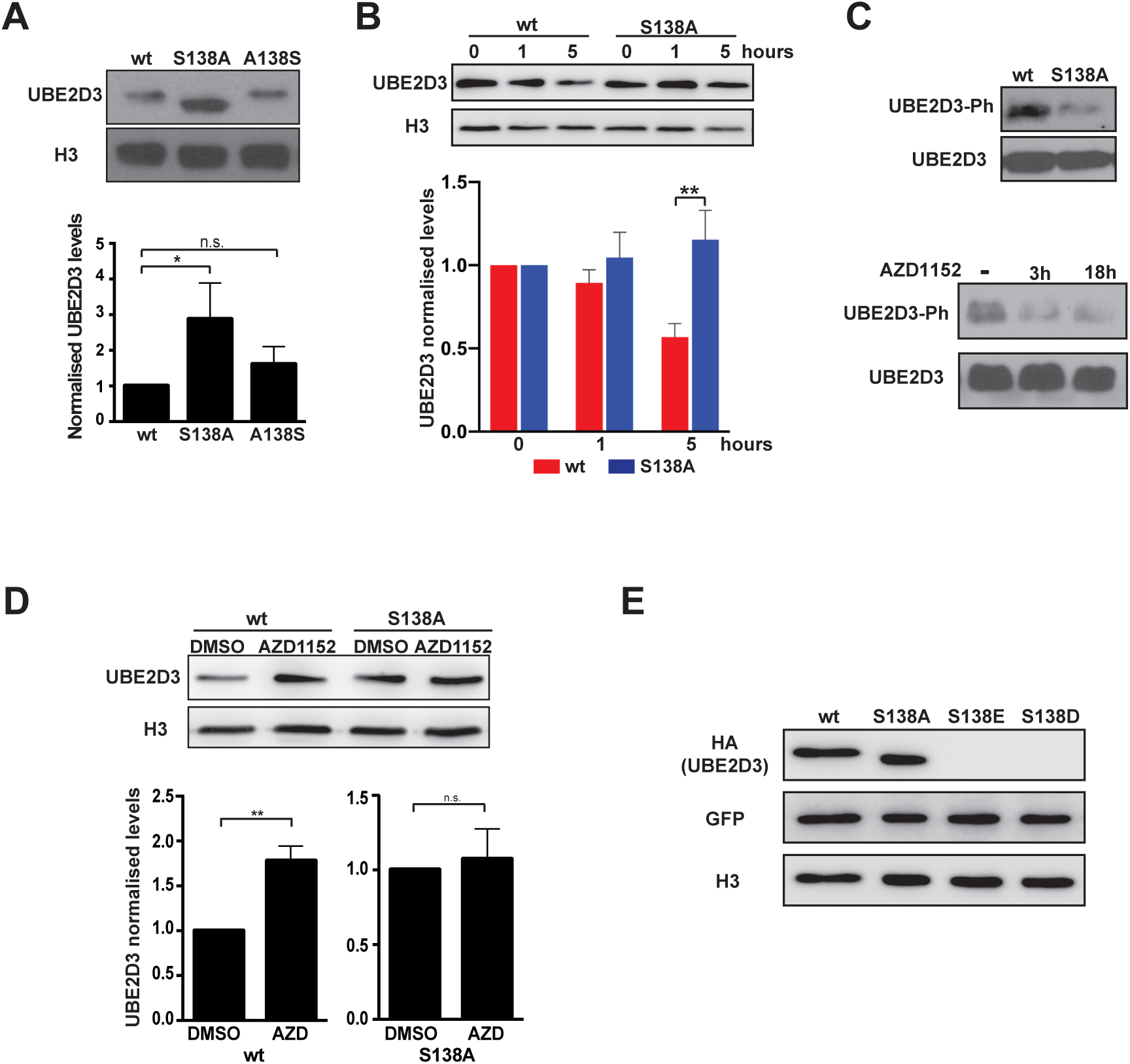
Characterization of UBE2D3 S138A mutant ESCs. (A) Western blot analysis of UBE2D3 levels in wt, S138A mutant and A138S revertant ESCs. Top panel: representative blot. Bottom panel: quantification of band intensity relative to histone H3; mean ± SEM, n = 5; p-values calculated by *t-*test for this, B and F: *p<0.05, **p<0.01. (B) Effect of protein synthesis inhibition in UBE2D3: western blot (top panel) and quantification (bottom panel) of UBE2D3 levels in wt and S138A mutant cells after treatment for 1 and 5 hours with 50 μg/μl cycloheximide. (C) Phosphorylation of UBE2D3-S138 in ESCs. Top panel: western blot of extracts from wt and UBE2D3 S138A ESCs with anti-phospho-UBE2D3 (UBE2D3-Ph) and anti-UBE2D3 (UBE2D3) antibodies. To allow direct comparison of the UBE2D3-Ph bands, samples were adjusted so that equivalent levels of total UBE2D3 were loaded. Bottom panel: western blot of extracts from wt ESCs treated with Aurora B inhibitor AZD1152 for 3 and 18 hrs and probed with anti-phospho-UBE2D3. (D) Western blot analysis of UBE2D3 levels in wt and S138A mutant ESCs treated for 3 hours with Aurora B inhibitor AZD1152 or vehicle (DMSO). PI staining and FACS analysis showed that AZD1152 had no effect on the cell cycle (data not shown). Top panel: representative blot. Bottom panel quantification relative to histone H3, mean ± SEM, n = 4. (E) Western blot analysis of wt, phosphomutant S138A and phosphomimetics S138E and S138D mutants expressed in ESC. Cells were sorted for expression of the IRES-GFP after the transient transfection.

### Phosphorylation of UBE2D3-S138 by Aurora B destabilises the protein in ESCs

Phosphorylation of UBE2D3-S138 (Frangini et al., 2013) has the potential to affect the stability of the protein by introducing a negative charge in the centre of the α4-helix of the protein. The presence of phosphorylated UBE2D3 in ESCs was demonstrated using an antibody that specifically recognises the phosphorylated UBE2D3-S138 residue (Frangini et al., 2013) (anti-UBE2D3-Ph; Figure S2D). Western blotting of protein extracts from wild-type and UBE2D3-S138A mutant ESCs using the anti-UBE2D3-Ph antibody gave a strong band in wild-type ESCs and only a low background band in the S138A mutant cells (Figure 2C, top panel). The background is likely to result from cross-reaction of the anti-UBE2D3-Ph antibody with the unphosphorylated protein that we have observed *in vitro*. Incubation of ESCs for 3 and 18 hours with the specific Aurora B inhibitor AZD1152 reduced the level of phosphorylated UBE2D3 to background levels (Figure 2C, bottom panel;), confirming the involvement of Aurora B in phosphorylating S138. These results show that Aurora B has a major role in phosphorylating wild-type UBE2D3-S138 in ESCs, although we cannot completely exclude the possibility that S138 is also a target for phosphorylation by other kinases.

To directly test whether phosphorylation of UBE2D3-S138 destabilises UBE2D3, wild-type and UBE2D3-S138A mutant ESCs were incubated for 3 hours with the specific Aurora B inhibitor AZD1152. The short incubation period was used to avoid any possibility of interference with the cell cycle. Western blot analysis showed that inhibition of Aurora B led to a significant increase in the level of UBE2D3 protein compared to cells treated with vehicle (Figure 2D). In contrast, no change was observed in the level of the UBE2D3-S138A mutant protein, where the Aurora B phosphorylation site is not present. As a further test of the effect of phosphorylation of S138, phosphomimetic mutant UBE2D3 proteins that had glutamic acid or aspartic acid residues at position 138 were transiently expressed in wild-type ESCs. The results show that the phosphomimetic mutants are very unstable in ESCs (Figure 2E), providing further confirmation of the destabilizing effect of phosphorylation of S138.

### UBE2D3-S138 phosphorylation alters the structure of the protein causing it to become insoluble and aggregate

To further investigate the effect of phosphorylation on UBE2D3, the wild-type, S138A and phosphomimetic S138E and S138D proteins were expressed in bacteria with the aim of purifying the proteins for structural analysis (see Methods). A western blot of the soluble fractions and the pelleted inclusion body fractions obtained after IPTG induction of expression followed by lysis of the bacteria is shown in Figure 3A. The bulk of the wild-type and S138A mutant UBE2D3 proteins was located in the soluble fractions whereas the phosphomimetic mutant proteins were almost entirely localised to the insoluble inclusion bodies, providing direct evidence that the presence of a negatively charged residue at position 138 disrupts the structure of UBE2D3. This observation also indicates that the C-terminal α–helix has an important role in regulating the stability and level of the protein.

**Figure 3.**
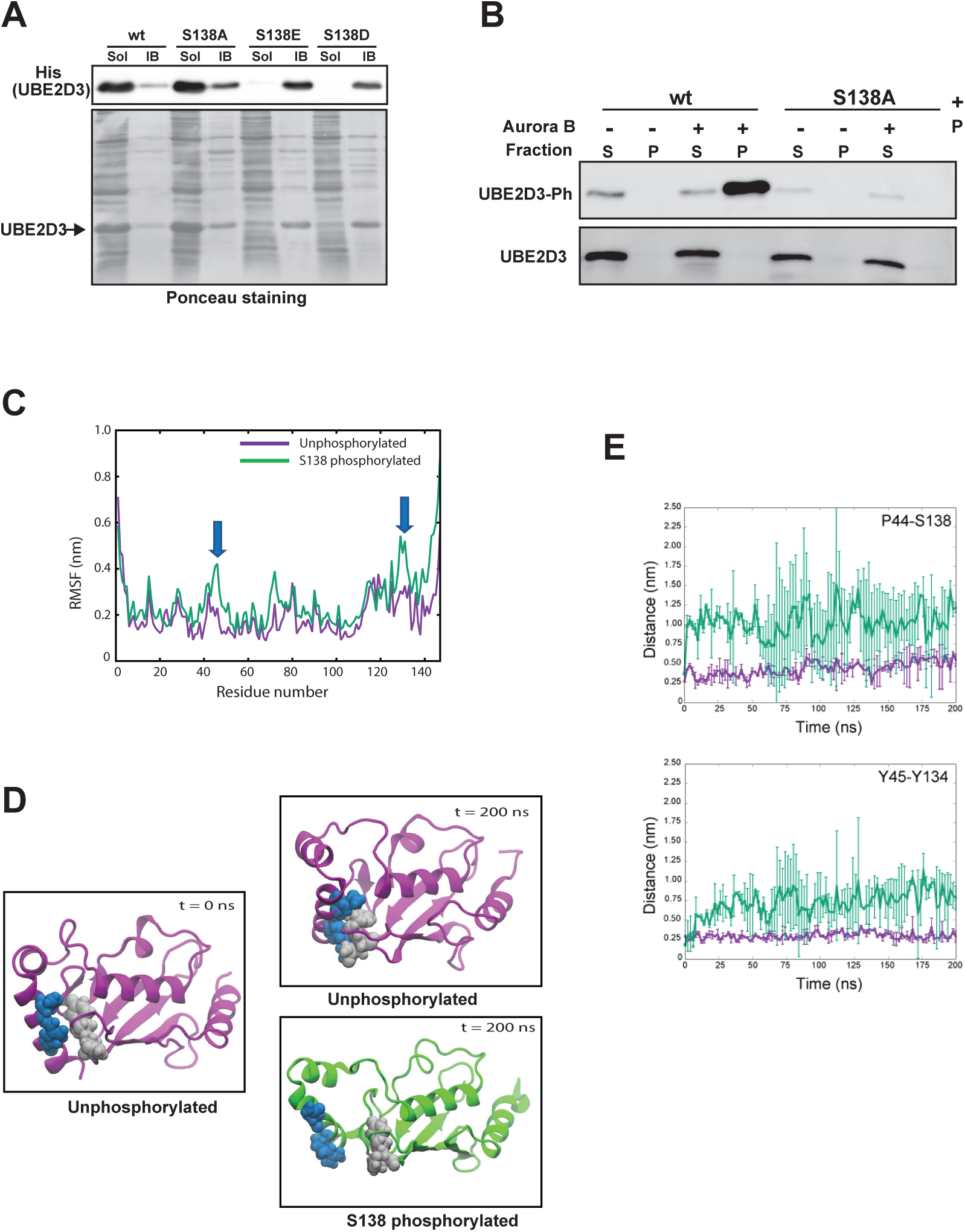
Effect of S138 phosphorylation on the structure of UBE2D3. (A) Solubility of UBE2D3 wt, phosphomutant S138A and phosphomimetic S138E and S138D variants. UBE2D3 was expressed in bacteria and extracted as described in methods. Soluble (Sol) and insoluble inclusion body (IB) fractions were analysed by western blot and probed with anti-His antibody. Ponceau Red staining was used as loading control. (B) Phosphorylation of UBE2D3 at S138 causes precipitation. Western blot showing bacterially expressed and purified wt and S138A proteins after *in vitro* phosphorylation and centrifugation, S = supernatant, P= pellet. (C – E) Results of molecular dynamics simulation of the effects of S138 phosphorylation on the structure of UBE2D3. (C) Root mean square fluctuation (RMSF) calculations for unphosphorylated and S138 phosphorylated UBE2D3. Blue arrows indicate regions of increased mobility in the C-terminal α4-helix where S138 is located and around P44. (D) Effect of S138 phosphorylation on the positioning of the C-terminal α4-helix. P44 and Y45 are shown in grey, Y134 and S138 in blue. The unphosphorylated protein is purple, and the S138 phosphorylated protein is green. In the S138 phosphorylated protein, the α4-helix has moved away from the loop carrying Pro44 and Tyr45, exposing the hydrophobic core of the protein. (E) Distance between key residues in the simulations of the wild type and phosphorylated proteins. The distances were measured between P44 and S138 (left panel) and between Y45 and Y134 (right panel) Each curve is averaged over 2 independent simulations. The bars indicate the standard deviations.

Efforts to purify soluble UBE2D3-S138E or S138D proteins by varying the bacterial growth conditions were unsuccessful, precluding direct functional analysis of the phosphomimetic mutant proteins. Therefore, as an alternative strategy, purified wild-type and UBE2D3-S138A proteins were incubated with Aurora B kinase and ATP. This was followed by centrifugation of the reaction mix and analysis of the supernatant and pelleted fractions by western blotting and probing with the anti-UBE2D3-S138Ph antibody. The results show clearly that the phosphorylated form of wild-type UBE2D3 is located almost entirely in the pelleted fraction (Figure 3B). No phosphorylation signal above background was observed in the pelleted or supernatant fractions for the UBE2D3-S138A mutant protein. Together with the observation that the phosphomimetic mutant proteins are insoluble when expressed in bacteria, these results provide strong evidence that phosphorylation of UBE2D3-S138 disrupts the structure of the protein, causing it to aggregate and become insoluble.

The effect of S138 phosphorylation on the molecular structure of UBE2D3 was also addressed by carrying out molecular dynamic simulations on the protein in the unphosphorylated and phosphorylated states (Figure 3C-E; see Methods). The results show that phosphorylation substantially alters the structure of the protein. Two independent simulations each of duration 200 ns were performed for the wildtype and phosphorylated protein. While both proteins remained stable in terms of secondary structure on the timescale of the simulations, distinct differences in their conformational behavior were observed. The unphosphorylated protein displayed much less flexibility compared to the phosphorylated protein. This is clearly evident from the root mean square fluctuation (RMSF) which measures the deviations from the average structure calculated over the simulation (Figure 3C). In particular residues 43-55, 70-80 and 130-138 show a marked increase in flexibility in the phosphorylated compared to the unphosphorylated protein.

Visual inspection of the trajectories (Figure 3D) revealed that upon phosphorylation, a distinct conformational rearrangement is observed in the phosphorylated protein, namely that the α4-helix, which contains the phosphorylated S138 moves away from the loop carrying Pro44 and Tyr 45, thereby exposing the hydrophobic core of the protein. This was further confirmed by calculating the distances between residues P44 and S138 and between Y45 and Y134, which were found to increase in the phosphorylated protein (Figure 3E). These results suggest that upon extension of the simulations, the phosphorylated protein is likely to begin to unfold. This rearrangement was not observed in the unphosphorylated protein, in which the α4-helix and the aforementioned loop remain in close proximity throughout the simulations.

### The UBE2D3-S138A mutation is an early embryonic lethal in mice

The evidence that phosphorylation downregulates the level of UBE2D3 in ESCs by disrupting the structure of the protein led us to consider the possibility that S138 phosphorylation could have acquired functional roles in embryogenesis during amniote evolution. To investigate the possible involvement of UBE2D3-S138 phosphorylation in amniote embryogenesis, we used the mouse embryo as a model system. The CRISPR/Cas9 targeting strategy described above was used to mutate S138 to the anamniote alanine in fertilized mouse eggs (see Methods). Initially, the eggs were implanted into foster mothers and allowed to develop to term. However, this did not yield any live mice that carried the mutation, which suggested that the S138A mutation is an embryonic lethal in mice. To determine whether this is the case, embryos generated by injection of fertilized eggs were dissected at 12.5 days after injection and re-implantation of the eggs (E12.5) and 5 tissues were dissected from each embryo and analysed by PCR to genotype the embryos and assess mosaicism (see Methods). The percentage of embryos carrying the S138A mutation at E12.5 was compared with the rate obtained by injection of a targeting sequence that was designed to generate a synonymous mutation (S138S) that altered the codon sequence, leaving the protein sequence unchanged (see Methods). Injection of the control S138S targeting sequence gave a mutation rate of 22% (10/45). In contrast, when the S138A targeting sequence was injected, 2% of E12.5 embryos (3/137) were positive for the mutation (Figure 4A). One of the 3 positive embryos obtained for the S138A mutation was dead and sequencing of the 2 live embryos showed evidence of additional mutations around the splice site upstream from the mutated codon 138 that could have compensated for the S138A mutation by downregulating expression of the UBE2D3 gene (data not shown). As CRISPR/Cas9 targeting is known to occasionally introduce unwanted mutations close to the targeting site, the observation that a small proportion of embryos were able escape from the effects of a gain of function mutation due to acquisition of additional compensating mutations was not unexpected. The 10-fold reduction in the proportion of E12.5 embryos carrying the S138A mutation compared with the targeting efficiency for S138S provides clear evidence that the S138A mutation has a high penetrance embryonic lethal effect in mice that affects the viability of the early embryo.

**Figure 4.**
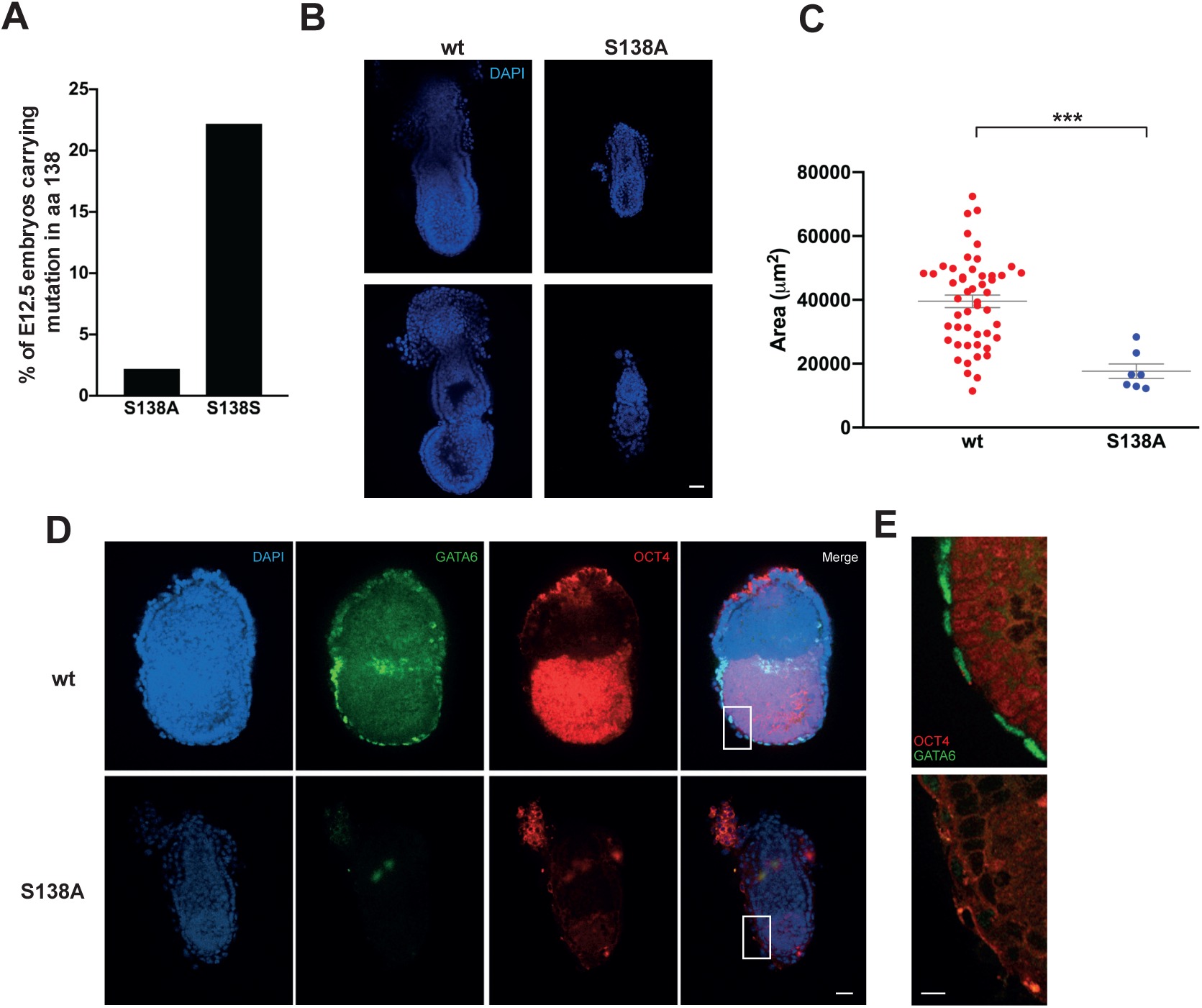
Early embryonic lethality of the UBE2D3-S138A mutation in mice. (A) Comparison of UBE2D3-S138A and -S138S mutation efficiency in E12.5 embryos obtained by CRISPR/Cas9 injection. (B) Representative wt and mutant E6.5 embryos. Scale bar = 50 μm. (C) Comparison of sizes for wt (n=50) and S138A (n=7) E6.5 embryos. The middle plane of each embryo was chosen for measurement; p-value was calculated by Mann-Whitney U test (***p<0.0001). (D) Wt and S138A E6.5 embryos stained with anti-OCT4 (epiblast) and anti-GATA6 (visceral endoderm). Scale bar = 30 μm. (E) Higher magnification of boxed region in panel F. Scale bar = 5 μm.

To further investigate the timing of the lethal effect, embryos were dissected 6.5 days after injection of the S138A targeting construct (E6.5 and E7.5) and analysed by microscopy. This was followed by PCR analysis to determine genotype (see Methods for details). Seven homozygous embryos were identified at E6.5 and confirmed by sequencing (Figure S3). A significant reduction in size was observed in the homozygous mutant S138A embryos compared with the wild-type embryos (p < 0.001) (Figure 4B, C). Embryos were also stained with anti-GATA6 and anti-OCT4 antibodies to assess visceral endoderm (VE) and epiblast development. Staining of the homozygous mutant E6.5 embryos gave no staining for GATA6 (VE) and very weak staining for OCT4 (epiblast) compared to wild-type cells (Figure 4D, E) The close up view of the peripheral region of the embryo shown in Figure 4E reveals the absence of the GATA6 staining that is indicative of VE cells at this stage of development. These results suggest that the presence of alanine at position 138 of UBE2D3 interferes with epiblast expansion and differentiation of VE in mouse embryos between E4.5 and E6.5.

In summary, the CRISPR/Cas9 mutagenesis in mice shows that mutation of UBE2D3-S138 to the alanine that is conserved in non-amniote eukaryotes is a high penetrance early embryonic lethal mutation in mice with death most likely caused by a failure of lineage commitment and cellular expansion between E4.5 and E7.5. These results support the idea that phosphorylation of UBE2D3-S138 acquired important functional roles in embryogenesis during amniote evolution.

### Phosphorylation of UBE2D3-S138A affects differentiation of embryonic primitive endoderm cells

The CBL E3 ubiquitin ligase plays an important role in regulating receptor tyrosine kinase (RTK) turnover by monoubiquitinating phosphorylated RTKs and promoting trafficking of the receptors to the lysosome for degradation. RTKs that are known targets for CBL include PDGFRα, EGFR and FGFR, all of which have essential roles in embryonic development. In particular, FGFR1 and PDGFRα are involved in differentiation and survival of primitive endoderm (PrE), which is a precursor of extraembryonic lineages. (Bessonnard et al., 2019; Molotkov et al., 2017) CBL has been shown to use UBE2D3 as a ubiquitin donor (Fortian et al., 2015; Liyasova et al., 2019), leading us to speculate that phosphorylation of UBE2D3-S138 mutant might be affecting the level of RTKs in embryonic cells by regulating the activity of CBL.

To test this possibility we induced wild-type and mutant ESCs to differentiate into XEN cells, which are an *in vitro* model for primitive endoderm (PrE), by incubating them with all-trans retinoic acid (ATRA) and activin (Niakan et al., 2013; Tomaz et al., 2017). PrE differentiates from the inner cell mass (ICM) of the developing blastocyst in response to signalling through the FGFR and PDGFRα RTKs (Bessonnard et al., 2019; Molotkov et al., 2017). The cells of the PrE lineage give rise to the parietal endoderm, which forms the inner surface of the yolk sac. PrE cells are also precursors of visceral endoderm, which has important inductive roles during gastrulation. After 8 days of induction, wild-type and revertant ES cells gave rise to large numbers of cells with the characteristic XEN cell morphology (Niakan et al., 2013; Tomaz et al., 2017), whereas the mutant cells failed to differentiate efficiently and gave significantly reduced numbers of XEN cells. (Figure 5A). The reduced efficiency of differentiation was confirmed by a time-lapse analysis from days 4 to 8 of the differentiation, which showed greatly reduced rates of XEN cell colony formation during differentiation of the S138A mutant cells compared with the wild-type cells. (Supplementary Video 1). FACS analysis of PDGFRα and immunofluorescent staining for PDGFRα and FGFR1 showed strong reductions in the levels of both RTKs in the mutant cells after 8 days of differentiation, confirming the impact of the UBE2D3-S138A mutation on PrE differentiation. (Figure 5B and 5C).

**Figure 5.**
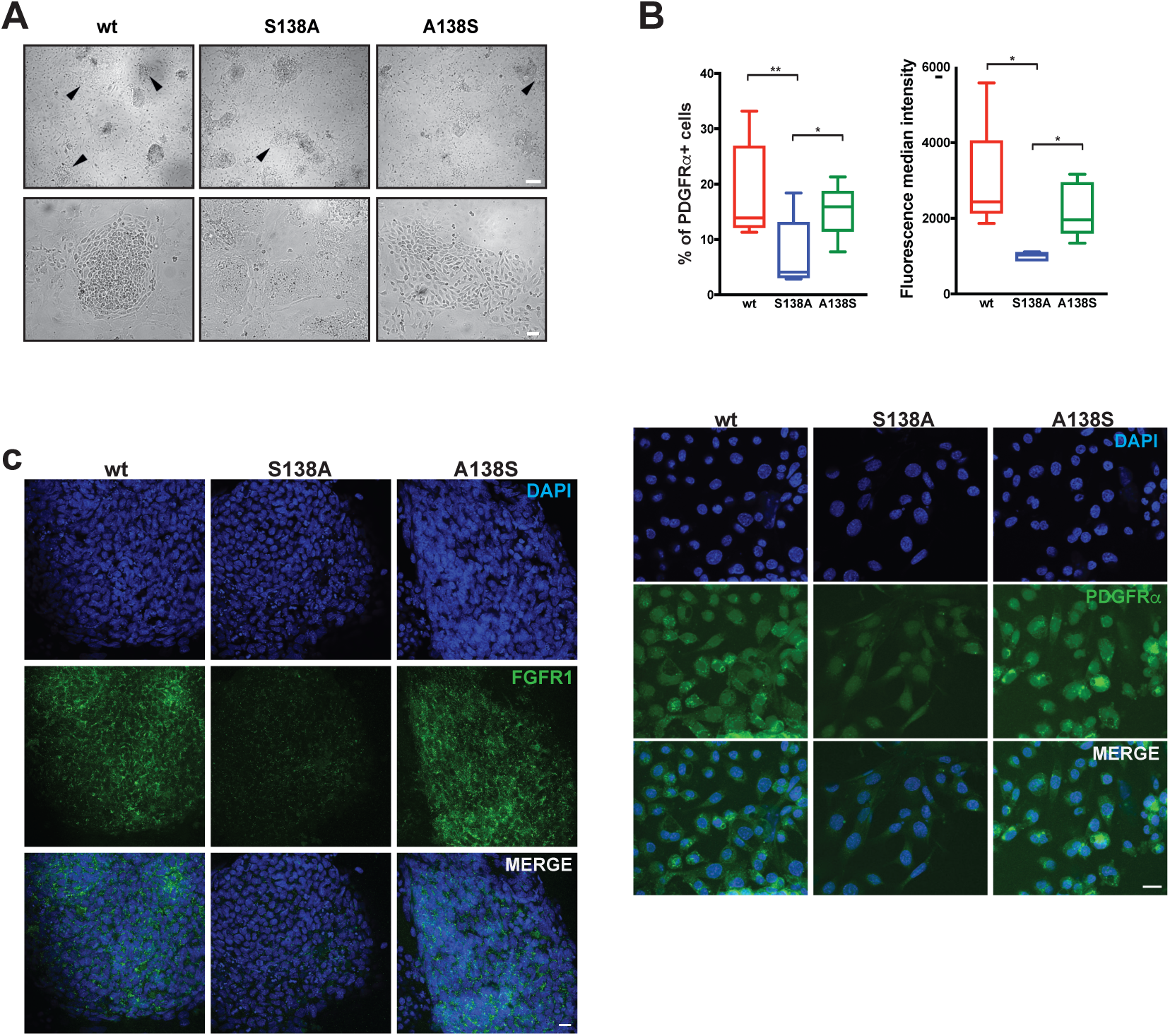
Impaired differentiation of UBE2D3-S138A mutant ESCs into XEN cells. (A) ESC differentiation into XEN cells. Bright-field images of wt, S138A mutant and A138S revertant cells at day 8 of differentiation; arrows indicate colonies with XEN cell morphology. Top panel scale bar = 500 μm, bottom panel scale bar = 100 μm. (B) Cells collected at day 8 of XEN cell differentiation were stained with anti-PDGFRα and analysed by FACS and immunostaining. Top left panel: percentage of cells that stained positive for PDGFRα. Top right panel: Measurement of fluorescence median intensity of staining of cells that are positive for PDGFRα (p-values calculated by *t*-test: *p<0.05, **p<0.01, n=5). Bottom panel: immunostaining of wt, S183A and revertant cells with anti-PDGFRα. Scale bar = 20 μm. (C) Immunostaining of wt, S183A and revertant cells with anti-FGFR1 on day 8 of differentiation to XEN cells. Scale bar = 20 μm.

### Interactions between CBL and its receptor tyrosine kinase substrates are enhanced by the UBE2D3-S138A mutation

The proximity ligation assay (PLA) was used to measure the interaction between CBL and UBE2D3 during differentiation of ESCs into XEN (primitive endoderm) cells (Figure 6A – D). PLA detects and amplifies interactions between oligonucleotide tags on the two antibodies, generating interaction foci that can be visualised by microscopy. When PLA was used to compare the level of interaction between UBE2D3 and CBL in wild-type, S138A and revertant A138S ESCs after 3 days of differentiation, a substantial increase in the number of interaction foci was observed in the S138A mutant cells compared with wild-type cells, whereas the A138S revertant cells showed a similar level of interaction to the wild-type cells (Figure 6B). These results provide evidence that phosphorylation of UBE2D3-S138 downregulates CBL activity during PrE differentiation and that this regulation is blocked by the S138A mutation.

**Figure 6.**
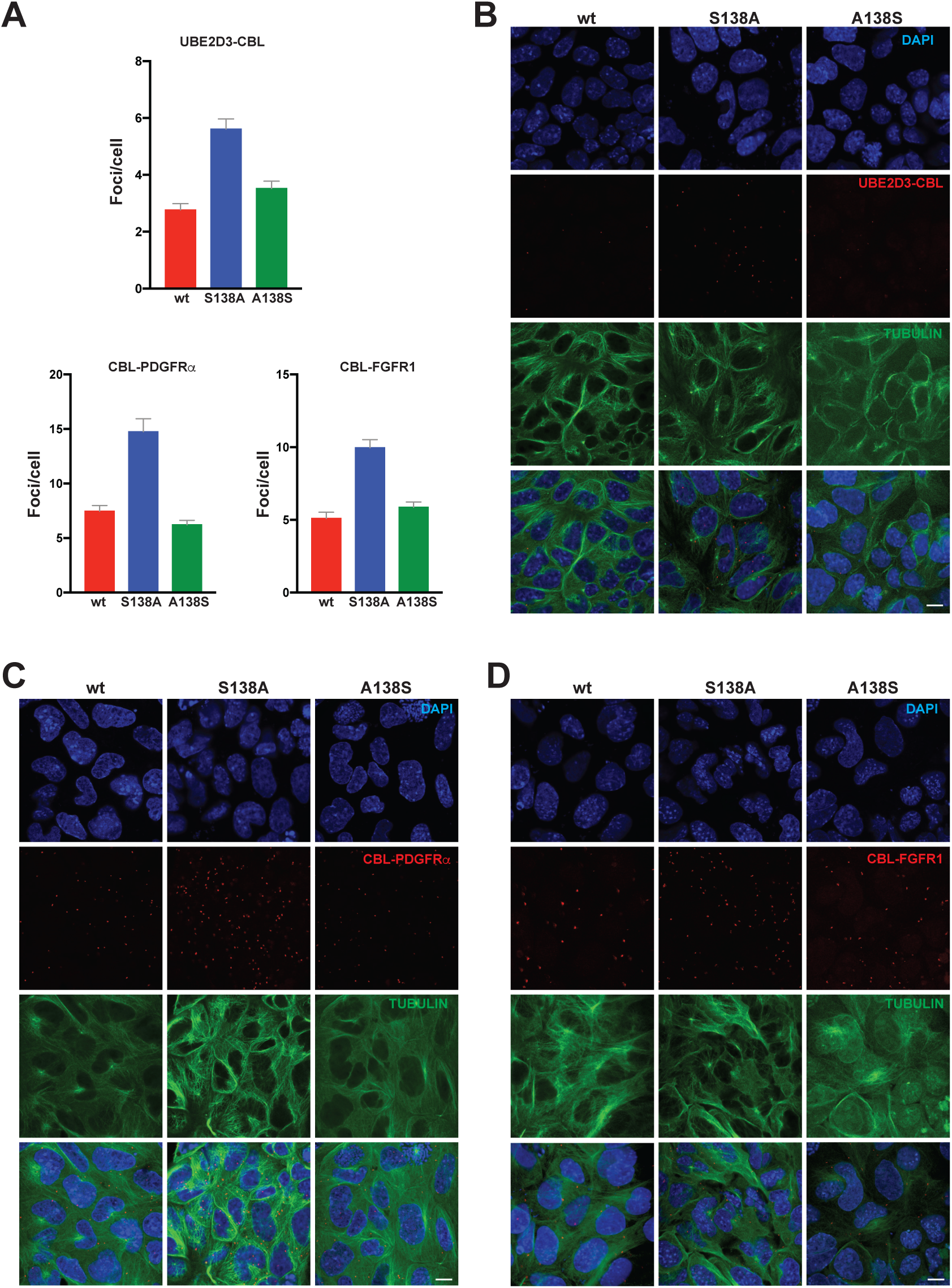
PLA of UBE2D3-CBL, CBL-PDGFRα and CBL/FGFR1 contacts at day 3 of XEN differentiation. (A) Quantification of the number of PLA foci/cell, all graphs represent mean ± SEM of 3 different experiments. (B-D) Representative images of PLA detected interaction between UBE2D3-CBL (B), CBL-PDGFRα (C) and CBL-FGFR1 (D).

PLA was also used to analyse the interaction between CBL and the RTK PDGFRα after 3 days of XEN cell differentiation. The results showed an increase in the number of interaction foci in S138A mutant cells compared with wild-types cells (Figure 6C). A similar increase was also observed in the interactions between CBL and FGFR1 (Figure 6D). The increased interactions provide an explanation for the reduction in the levels of PDGFRα and FGFR1 observed by immunofluorescence staining in differentiating PrE cells.

The effect of the S138A mutation on the ability of ESCs to differentiate into the three germ layers, ectoderm, mesoderm and endoderm, was also tested (See Methods). Differentiation into the germ layers showed that wild-type and mutant ESCs gave rise to all three cell types with similar efficiency (Figure S4). These results show that the effect of Aurora B downregulation of UBE2D3 levels and CBL activity is strongest on differentiation of extraembryonic PrE, although we cannot rule out more subtle effects of the mutation on differentiation of the germ layers that would contribute to the lethal effect of the UBE2D3-S138A mutation in mouse embryos.

UBE2D3 acts a ubiquitin donor for a number of E3s in addition to CBL and is involved in the ubiquitination of a wide range of substrates. Cyclin D1 is a known target of UBE2D3-mediated ubiquitination, which leads to its downregulation (Hattori et al., 2007; Mittal et al., 2011). Analysis of Cyclin D1 levels in the analysed in the S138A mutant ESCs showed that the level was significantly reduced in the mutant cells (Figure S2E). As a further test of the broad effects of the S138A mutation, the level of monoubiquitination of histone H2AK119 (H2AK119Ub) was measured at gene promoters in ESCs. H2AK119 ubiquitination is mediated the E3 ligase activity of the Polycomb protein RING1B using UBE2D3 as a ubiquitin donor, and has been shown to be important for maintaining poising of developmentally regulated promoters in ESCs (Stock et al., 2007). Chromatin immunoprecipitation (ChIP) analysis was carried out on chromatin isolated from wild-type and mutant ESCs using primers for a range of silent and active promoters. The results show a clear increase in the level of H2AK119Ub across the majority of promoters analysed (Figure S5A and S5B) and a reduction in binding of S5-phosphorylated RNA polymerase II (RNA Pol II-S5Ph) (Figure S5C). Interestingly, we observed increased H2AK119Ub and reduced RNA Pol II-S5Ph at active as well as silent promoters. This is consistent with a previous report that RING1B binds to and ubiquitinates active as well as silent promoter regions in ESCs (Brookes et al., 2012). Overall, our results show that the increased UBE2D3 levels caused enhanced ubiquitination by a number of different E3s. This highlights the broad effect that phosphorylation of UBE2D3-S138 has on ubiquitination and provides a potential explanation for the very high penetrance phenotype that was observed for the S138A mutation in mouse embryos.

## DISCUSSION

Amniotes first appeared during the Carboniferous period between 314 and 340 Mya (Carroll, 1964; Clack, 2012) and ultimately became the dominant vertebrate life forms on dry land. The results of this study identify an alanine to serine substitution that occurred in the common ancestor to modern amniotes, generating a novel mechanism for regulating the promiscuous ubiquitin-conjugating enzyme UBE2D3. The conservation of the alanine residue at position 138 in UBE2D3 from single celled eukaryotes through to modern invertebrate metazoan and anamniote vertebrates, together with the absolute conservation of the serine residue in amniotes provides strong evidence that the substitution was a highly unusual event that happened only once in eukaryotic evolution. Acquisition of the serine created a phosphorylation site for the cell cycle kinase Aurora B, and we have shown that phosphorylation of this site drastically reduces the stability of the UBE2D3 protein by altering the position of the C-terminal α-helix and exposing the hydrophobic core of the protein. This causes a reduction in the solubility of the protein, targeting it for degradation.

Experiments in mice and ESCs showed that phosphorylation of UBE2D3-S138 is required for mouse embryonic development. Intriguingly, the strongest effect of the mutation that we observed in ESCs and embryos was on the differentiation of extraembryonic endoderm lineages (PrE and VE), which are known to have important roles in the formation of the amniote extraembryonic membranes and in the positioning of the primitive streak during gastrulation in chick and mouse embryos (Ferner and Mess, 2011; Stern and Downs, 2012).

The experiments using PLA provide clear evidence that the S138A mutation affects the functioning of the E3 ligase CBL, with increased interaction between UBE2D3 and CBL and between CBL and two of its target RTKs, PDGFRα and FGFR1, observed during differentiation of S138A mutant ESCs into XEN cells. Ubiquitination by CBL is essential for mouse development, with mice that are null for the two major isoforms c-CBL and CBL-b showing an embryonic lethal phenotype before E10.5 (Naramura et al., 2002). Our results provide evidence that regulation of CBL activity is also essential for mouse embryonic development and we show that the increased CBL activity resulting from upregulation of UBE2D3 in the S138A mutant cells is associated with downregulation of FGFR1 and PDGFRα in differentiating XEN (PrE) cells. In addition to their roles in PrE differentiation, FGFR1 is involved in embryonic growth and mesodermal patterning during gastrulation (Deng et al., 1994; Yamaguchi et al., 1994) and PDGFRα is required for regulation of neural crest development, somite patterning and craniofacial development (Soriano, 1997; Xu et al., 2005). Reduced activity of both of these RTKs is likely to contribute to the very strong effect embryonic lethal effect that we observe for the UBE2D3-S138A mutation.

In addition to the effects on CBL, we also observed increased ubiquitination of histone H2AK119 at repressed and active genes in S138A mutant ESCs and downregulation of Cyclin D1. Both of these proteins are known targets for ubiquitination that uses UBE2D3 as a ubiquitin donor (Bentley et al., 2011; Mittal et al., 2011). It is notable that we did not obtain significant numbers of heterozygotes or mosaics for the S138A mutation at E12.5, which suggests that the mutation causes a general cell autonomous defect that affects survival of the majority of cell lineages in the developing embryo. This could be a reflection of the number of different E3s that are able to use UBE2D3 as a ubiquitin donor.

The promiscuous activity of UBE2D3 is an intrinsic function of the protein (Brzovic and Klevit, 2006) that is highly conserved, as evidenced by the very high homology exhibited by UBE2D3 orthologues across eukaryotic lineages. UBE2D3 also has the ability to synthesise different types of ubiquitin chains. In addition, UBE2D3 operates in a complex regulatory environment within the cell, which is populated in mammals by around 30 E2 enzymes with varying specificities for around 600 partner E3s. The functioning of E2 enzymes involves co-operative action between different E2s and also, potentially, competition between E2s for access to partner E3s. Promiscuous proteins are known to confer evolvability by virtue of being able to adopt novel functions and regulatory roles (Aharoni et al., 2005). We hypothesise that the A138S substitution and the acquisition of phosphoregulation of UBE2D3 levels by Aurora B at the anamniote to amniote transition is an example of this type of acquisition that altered the ubiquitination landscape in the ancestral amniote lineage.

## MATERIAL AND METHODS

### Comparative analysis of UBE2D3 amino acid sequences

#### Vertebrate sequences

The UBE2D3 protein sequence comparison was based on protein sequences downloaded from the UCSC 100-way MULTIZ alignment for hg19 exons against 100 vertebrate species. Protein sequences were checked manually for accuracy using gene sequences from the UCSC browser, NCBI or ENSEMBL (see below). UBE2D3 genes were identified by the presence of an additional minor alternative exon (exon-7a) that lies upstream from the predominantly used exon-7b and is specific to UBE2D3 (see Supplementary Methods). Manually annotated fish UBE2D3 sequences, in particular, differed significantly from the sequences produced by automated annotation. This is probably due to the high homology between UBE2D3 family members. Sources and sequence co-ordinates or accession numbers for the UBE2D3 gene are shown in Table S2. Full details of the approaches used to identify UBE2D3 genes and verify the protein sequences can be found in Supplementary Methods.

#### Carolina Anole Lizard (*Anolis carolinensis*)

Due to problems with annotation of the UBE2D3 sequence of the Carolina Anole Lizard, the sequence was obtained by mining of transcriptome data (see Supplementary Methods).

#### Axolotl (*Ambystoma mexicanum*)

Candidate partial UBE2D3 sequences from were obtained by blasting the Axolotl genomic sequence using the exon-6 and exon-7 sequences from the Tibetan frog (*Nanorana parkeri*). As it was not possible to determine which of these sequences is UBE2D3 due to the absence of the diagnostic exon-7a, both are shown in Figure 1 as Axolotl-1 and −2 (see Supplementary Methods).

#### Giant Panda (*Ailuropoda melanoleuca*)

The annotated UBE2D3 sequence for the Giant Panda contains a cluster of 4 variant residues at positions 112, 113, 115, 116. Since 3 of these residues are invariant in all other eukaryotes examined, we concluded that the apparent variation in the Giant Panda sequence was the result of a sequencing error. Therefore, the Giant Panda was omitted from the sequence comparison.

#### Sequences from non-vertebrate eukaryotes

Sequences of UBE2D3 homologues from representatives of non-vertebrate eukaryotic lineages were mined from the Uniprot database by blasting the entire database with conserved sequences from amino-terminal and C-terminal regions of *Drosophila melanogaster* UBCD1 that did not include position-138. The sequences used for the blast were: MALKRINKEL; REWTRKYA; REWTRKYAM. Sequences were also obtained by examination of protein sequences of individual E2 conjugating enzymes from model organisms in the Uniprot database.

### Mice

All work involving mice was carried out under the regulations of the British Home Office and was approved by the Imperial College Animal Welfare and Ethical Review Body. To generate mouse embryos with the UBE2D3-S138A mutation, gRNA for the mutation was transcribed *in vitro* and injected in C56Bl6/CBA F1 fertilized oocytes together with Cas9 mRNA and the donor ssDNA. To obtain E6.5, E7.5 and E12.5 embryos, the injected embryos were transplanted into foster mothers and harvested at the appropriate time point.

### Cells

E14 ESCs (female, 129/Ola) were obtained from the LMS Stem Cells and Transgenics facility. E14 ESCs were maintained on gelatin coated plates, at 37°C, 5% CO_2_, in KnockOut Dulbecco’s Modified Eagle’s Medium supplemented with 15% fetal bovine serum, 1x NEAA, 2 mM L-glutamine, 100 units/ml penicillin, 100 μg/ml streptomycin, 100 μM βME (all reagents from ThermoFisher) and 1000 units/ml LIF (Merck).

#### ESC differentiation into XEN cells

Differentiation was performed using the protocol detailed by (Niakan, et al., 2013). For time-lapse imaging of differentiation (see Supplemental Video 1), cells were transferred on day 4 of differentiation to a Zeiss Axiovert 200 microscope (Zeiss) where images were taken at 10 minute intervals until day 8. Images were analysed using Volocity software (Perkin Elmer).

#### Aurora B inhibition

ESCs were incubated for 3 hours with 200 nM AZD1152 (Merck) or DMSO in medium.

#### Protein synthesis inhibition

ESC were incubated for 1h and 5h with 50 μg/ml of cycloheximide (Cell Signalling Technology) or DMSO in medium.

### UBE2D3 gene editing using CRISPR/Cas9

The guide RNA (gRNA) used to generate the Serine to Alanine mutation was 5’-CACCGCAGGTACAACAGAATATCTC-3’ and the gRNA used to revert the mutation was 5’-CACCGACAACAGAATAGCCCGCGAA-3’. The gRNAs were cloned into pX330 vector (Cong *et al.*, 2013; Addgene plasmid #42230).

The donor ssDNAs used in this study are: 5’-TTAATTTTATTTTTTAAATAGCTTATTTGTTTGTTTACAGGTACAACAGAATAGCCCGCGAATGGACTCAGAAGTATGCCATGTGATGCTACCTTACAGTCAGAATAACC-3’ carrying the S138A mutation; 5’-TTAATTTTATTTTTTAAATAGCTTATTTGTTTGTTTACAGGTACAACAGAATATCTCGCGAATGGACTCAGAAGTATGCCATGTGATGCTACCTTACAGTCAGAATAACC-3’ to re-establish the wild type sequence and 5’-TTAATTTTATTTTTTAAATAGCTTATTTGTTTGTTTACAGGTACAACAGAATAAGCCGCGAATGGACTCAGAAGTATGCCATGTGATGCTACCTTACAGTCAGAATAACC-3’ to introduce an S138S synonymous mutation.

To produce E14 UBE2D3-S138A mutant ESC, 4×10^6^ cells were transfected with 3 μg of pX330 plasmid carrying the appropriate gRNA, 4 μg of the donor ssDNA and 3 μg of a puromycin resistance plasmid (pCAG-puro^R^) using the Mouse ES Cell Nucleofector™ Kit (Lonza) following the manufacturer’s protocol. One day after transfection cells were subjected to puromycin selection (1.5 μg/ml) for 24h. A week after transfection, individual clones were picked and genotyped by nested PCR. Mutant genotypes were confirmed by sequencing.

### Flow cytometry

All experiments were performed in a LSRII (Becton-Dickinson) and analysed with FlowJo Software. Cells were harvested, washed twice with PBS and incubated at a densitiy of 10^6^ cell/ml with anti-PDGFRα (1:100) in 2%FBS-PBS for 20 min at 4°C. After incubation, cells were washed twice, resuspended in 2%FBS-PBS at 5×10^5^ cells/ml and analysed.

#### Cell cycle analysis

ESCs were trypsinised, washed twice with PBS, fixed with 70% ethanol for 30 min on ice and washed twice with 2% FBS-PBS. Finally, cells were resuspended in PBS containing 1μg/ml RNAse A (ThermoFisher), 50 μg/ml propidium iodide and 0.05% NP40, incubated 20 min at room temperature in the dark followed by analysis.

### Whole-mount immunostaining

Embryos were harvested at the appropriate time point and fixed in 4% paraformaldehyde + 0.01% TritonX100 + 0.1% Tween for 20 min at 4°C. Fixed embryos were washed 3 times for 5 min in 0.1% Triton-PBS (PBT) at room temperature. Embryos were then blocked and permeabilised in blocking solution (2% donkey serum-0.4% TritonX100-PBS) for 2 hours at room temperature. Incubation with the primary antibodies was carried out at 4°C for 2 days with the primary antibodies diluted in blocking solution. Embryos were then washed 3 x 15 min in PBT, incubated with the appropriate secondary antibodies diluted in blocking solution for 2 hours at room temperature, washed again 3 times 15 min in PBT and left ON in PBT+1μg/ml DAPI at 4°C until imaging the next day. Images were acquired in a Leica SP8 confocal microscope with LAS X software (Leica) and analysed using Fiji (Schindelin et al., 2012). The antibodies used for embryo staining and their dilutions are indicated in Table S4.

### Cell immunostaining

Cells were fixed with 4% PFA at 37°C during 15 minutes, after this time they were washed 3 x 5 min with PBS. Cells were then permeabilised with 0.4% Tx-100 in PBS for 5 min at room temperature and blocked with 10% donkey serum-0.1% Tx-100-PBS. Incubation with the primary antibodies was carried out overnight at 4°C. After the primary antibodies the cells were washed 3×5 min with PBS and incubated 1h at room temperature with the appropriate secondary antibodies. Cells were washed 3 x 5 min and incubated with 1μg/ml DAPI in PBS for 5 min. Images were acquired in a Leica SP8 confocal microscope with LAS X software (Leica) or in an Olympus IX70 microscope. Images were analysed with Fiji software. The antibodies used in this study and their dilutions are listed in Tables S6.

### Proximity Ligation Assay

PLAs were performed with Duolink in situ PLA (Sigma, mouse and rabbit probes) following the manufacturer’s instructions. In brief, cells were fixed with 4% PFA at 37°C during 15 min, washed 3×5 min with PBS, permeabilized with 0.4% Tx-100 in PBS for 5 min at room temperature and blocked 1 hour at 37°C with blocking solution. Incubation with the primary antibodies was performed at 4°C overnight (the antibodies used and their dilutions are indicated in Table S6). The following day, cells were washed 2×5 min with buffer A, incubated with the minus and plus probes for 1h at 37°C in antibody diluent solution, washed 2×5 min with buffer A, incubated with Duolink ligation mix 30 min at 37°C, washed 2×5 min with buffer A, incubated with Duolink amplification mix 1h and 40 min at 37°C, washed 2×10 min with buffer B, 1×5 min with buffer A and counterstained with anti-Tubulin FITC for 1h at room temperature. After counterstaining, cells were washed 2×2 min with buffer A, 1×1 min with 0.01x buffer B and incubated 5 min with 1μg/ml DAPI in PBS. All incubations were performed in a humidity chamber. Images were acquired in a Leica SP8 confocal microscope with LAS X software (Leica), PLA foci were quantified with Imaris software (Oxford Instruments).

### Transient expression of UBE2D3 mutant proteins in ESCs

Wt UBE2D3, S138A, S138E and S138D were cloned into the expression vector pCAG-IRES-GFP (Addgene, plasmid #11159). E14 cells (5 x 10^6^) were transfected with 10 μg of the appropriate plasmid with the Mouse ES Cell Nucleofector™ Kit (Lonza) and allowed to grow for 48h after transfection. Cells were then collected and sorted for GFP expression. UBE2D3 protein levels were analysed by western blot.

### Protein analysis by western blotting

Cells were lysed in ice cold RIPA buffer (50 mM TrisHCl pH 8, 150 mM NaCl, 1% NP40, 0.5% sodium deoxycholate, 0.1% SDS) with Complete™ protease inhibitor cocktail (Roche) for 10 min, sonicated (MSE Soniprep150, 3 times 5 sec on/1 min off, 14 μm amplitude) and cleared by centrifugation for 10 min at 14000g. Concentration of the extracts was measured with the BCA Protein Assay Kit (ThermoFisher) according to the manufacturer’s instructions. Extracts were diluted in 6x SDS loading buffer, boiled for 5 min and subjected to SDS PAGE. Gels were transferred to nitrocellulose membranes using a Bio-Rad wet tank blotting system. Membranes were blocked with 5% bovine serum albumin (BSA) in Tris-buffered saline with 0.05%NP40 (TBS-NP40) for 1h at room temperature and incubated with the primary antibody diluted in 0.5% BSA TBS-NP40 O/N at 4°C. On the following day, membranes were washed 3 times for 20 min with TBS-NP40, incubated with the appropriate secondary antibody for 1h at room temperature, washed again 3 times 15 min and developed using Millipore Crescendo ECL (Merck). For western blot quantification, membranes were developed and analysed using the ImageQuant LAS400mini and the ImageQuantTL Software (GE Healthcare). The antibodies used are listed in Table S6.

In the case of western blots probed with the anti-UBE2D3-S138Ph antibody, membranes were denatured prior to blocking by incubation for 30 min at 4°C with 6M guanidine hydrochloride, 20 mM Tris HCl pH 7.4, 1mM PMSF and 5 mM *β*ME solution.

### Protein expression in bacteria and extraction

The cDNAs of UBE2D3 variants (wt, S138A, S138E, S138D) were cloned into the pET-N-His (Origene) vector containing a 6xHis N-terminal tag. *E. coli* BL21(DE3) cells (Agilent) were transformed with the pET-UBE2D3 plasmids and protein expression was induced with 0.4 mM IPTG (Merck) for 3h at 37°C. Bacteria were pelleted and incubated in lysis buffer (50 mM Tris-HCl pH 8.0, 150 mM NaCl, 10 mM imidazole, 7 mM *β*-Mercaptoethanol, 300 μg/ml lysozyme) for 30 min at 4°C. Lysate was then sonicated and spun (15000g, 10 min, 4°C). Supernatant was collected as soluble fraction and pellet was subjected to the same lysis as before. The pellet obtained after the second lysis-sonication was resuspended in lysis buffer, sonicated one more time and collected as the insoluble fraction.

### *In vitro* phosphorylation assay

Recombinant His-UBE2D3 (1 μg) was incubated with 200 ng of GST-AuroraB (MRC PPU, College of Life Sciences, University of Dundee, Scotland, mrcppureagents.dundee.ac.uk) for 30 min at 30°C in phosphorylation buffer (50 mM TrisHCl pH 7.5, 10 mM MgCl_2_, 500 μM ATP, 1 mM DTT, 5 mM NaF). Reactions were stopped by addition of SDS loading buffer, boiled for 5 min or used for subsequent ubiquitination assays. Phosphorylation was analysed by western blot.

For the kinase and spin assay, in vitro phosphorylation reactions were performed as indicated above and the spun for 15 min at 15000g at 4°C, washed 2x with phosphorylation buffer and subsequently analysed by western blot.

### Molecular Modeling

Ubiquitin-conjugating enzyme E2 D3 (UBE2D3) structure was retrieved from the Protein Data Bank (PDB: 3UGB) (Page et al., 2012), along with the conjugated ubiquitin. Two systems were set up: one with the original wild-type structural and one with a phosphorylation on the Ser138 side-chain. Ubiquitin was maintained on both simulations, but excluded from the analysis of results. The phosphorylated Ser138 (Sep138) model was obtained by submitting the original structure to the CHARMM-GUI server (Jo et al., 2014; Jo et al., 2008). Parameters to describe the thioester bond between UBE2D3 and ubiquitin were generated by submitting a dipeptide composed by the two residues that participate in this interaction (Gly and Cys) to the CGenFF server (Vanommeslaeghe et al., 2010; Yu et al., 2012) and adapted into the CHARMM36 force field.

#### Atomistic Molecular Dynamics Simulations

Simulations were performed using the GROMACS simulation suite (2018.3 version) (Abraham et al., 2015), the CHARMM36 force field (Huang and MacKerell, 2013) and the TIP3P water model (Jorgensen et al., 1983). Simulations were performed for 200 ns each, with two independent replicates for each system (800 ns, overall). The chosen temperature was 340 K in order to enhance sampling and denaturation of the structures. It was maintained by making use of the velocity rescale thermostat (Bussi et al., 2007) with a time constant of 1ps. For pressure coupling, the Parrinello-Rahman barostat (Parinello et al., 1981) was employed to keep pressure at 1 atm with a time constant of 1 ps. Particle Mesh Ewald method (Darden et al., 1993) handled the long-range electrostatics. The LINCS algorithm (Hess et al., 2008) was applied to constrain bonds, allowing an integration step of 2 fs. Long-range electrostatic and van der Waals cut-offs were set at 1.4 nm. Charges were neutralized by the addition of counterions and an additional 0.15 M concentration of NaCl ions were included in the simulations. VMD (Humphrey et al., 1996) was employed for molecular visualization, and analyses were performed using GROMACS tools.

### Statistical Analysis

All statistical analyses were performed with GraphPad Prism software. The statistical tests used in each experiment and significances are indicated in the corresponding figure legends.

### Genotyping, ChIP-qPCR and RNA extraction

For details of these procedures, see supplementary methods.

## Acknowledgements

We thank, Dirk Dormann and Chad Whilding from the LMS Microcopy facility and Laurence Game and Ivan Andrews from the LMS genomics facility for experimental assistance, Elodie Ndjetehe for her help with CRISPR/Cas9 injections, Tom Carroll for assistance with the initial bioinformatics analysis and Tobias Warnecke for advice on sequence alignments. The work was supported by the LMS/NIHR Flow Facility at Imperial. This research was funded by the Medical Research Council UK.

---

**Supplementary Video 1.** Time lapse imaging of differentiations of wild-type (wt), UBE2D3-S138A mutant and A138S revertant ESCs into XEN cells. Images were taken at 10 minute intervals from day 4 to day 8 of the differentiation.

