## Supplementary data for "Evolution of an amniote-specific mechanism for modulating ubiquitin signalling via phosphoregulation of the E2 enzyme UBE2D3"

**Roman-Trufero et al.**

**Supplementary Information**

### Figure S1

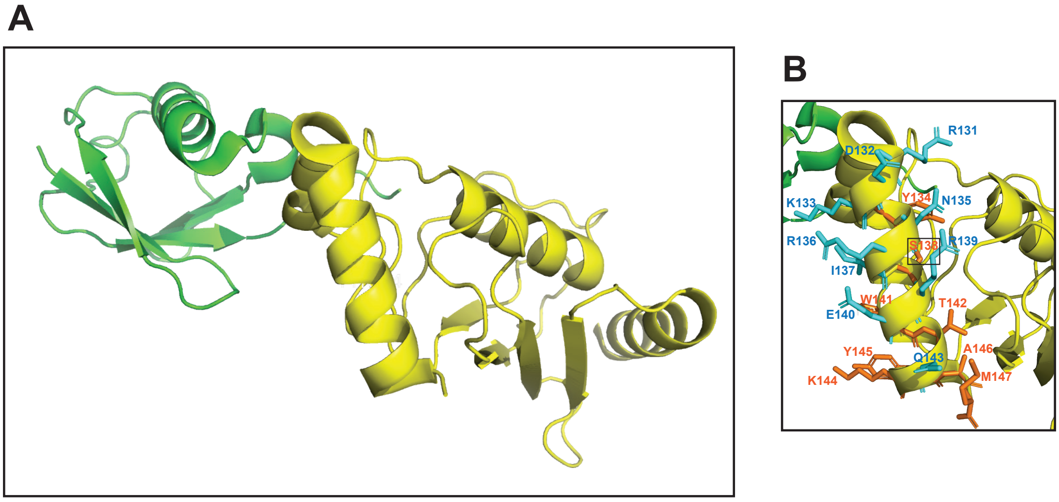


**Figure S1**. Structure of the human UBE2D3-ubiquitin conjugate. (A) UBE2D3 is shown in yellow and the conjugated ubiquitin molecule is coloured green. (B) Close up of the C-terminal α4-helix showing the side-chains of residues 131-147. Side-chains of variable residues are shown in cyan and side-chains of residues that are largely invariant are coloured orange. S138 (indicated by a box) is classified as largely invariant as it changes only once during evolution, at the anamniote/amniote transition. Position 138 is occupied by serine in amniotes and alanine in anamniotes and non-vertebrate eukaryotes. The images were prepared by visualising PDB 3UGB using PyMOL (v1.8.6.0.).

#### Figure S2

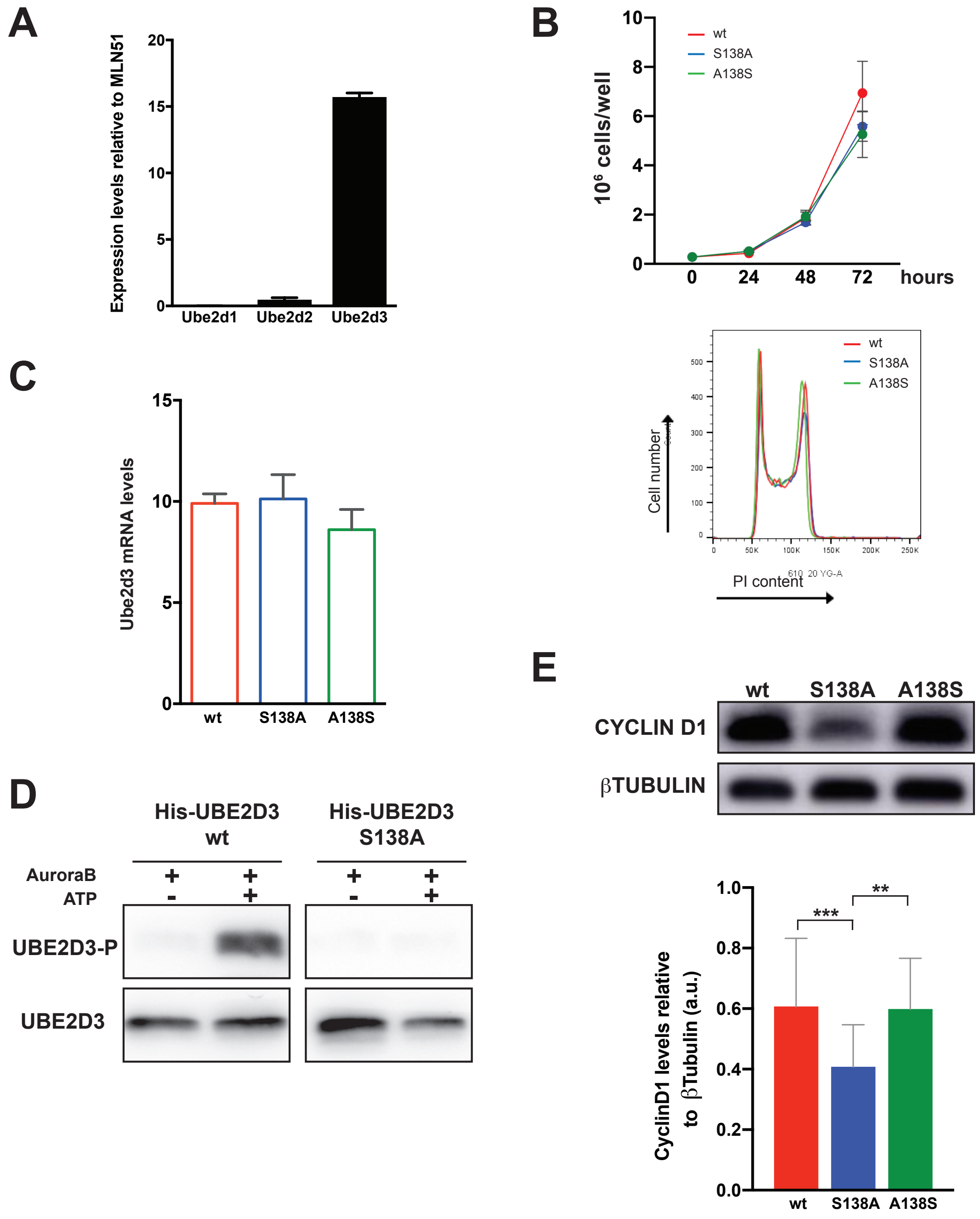


**Figure S2.** Characterisation of the UBE2D3 Ser138Ala mutant and Ala138Ser revertant ESC. (A) qRT-PCR analysis of UBE2D family members RNA in ESC. Transcripts were quantified using expression of the Mln51 gene as a standard (Szutorisz et al., 2005). (B) Top panel, proliferation assay: cells were plated at 10000 cells/cm$^2$ density and allow to grow for 3 days. Bottom panel: Analysis of cell cycle profiles; wild-type, mutant and revertant cells were stained with PI and analysed by FACS. (C) qRT-PCR analysis of UBE2D3 RNA in wild-type, Ser138Ala mutant and Ala138Ser revertant ESC. Transcripts were quantified as in (A). (D) Specificity of the anti-UBE2D3-S138p antibody. His-tagged UBE2D3 and UBE2D3-Ser138Ala were expressed in bacteria, purified and incubated with Aurora B kinase and ATP. Western blotting was carried with anti-UBE2D3-P (upper panels) and anti-UBE2D3 (lower panels). (E) Western blot analysis of CyclinD1 expression in wt, S138A and A138S. Top panel: representative western blot. Bottom panel: quantification relative to βTubulin.

All graphs represent mean ± SEM; (A), (B) and (C) n= 3; (E) n = 8.

#### Figure S3

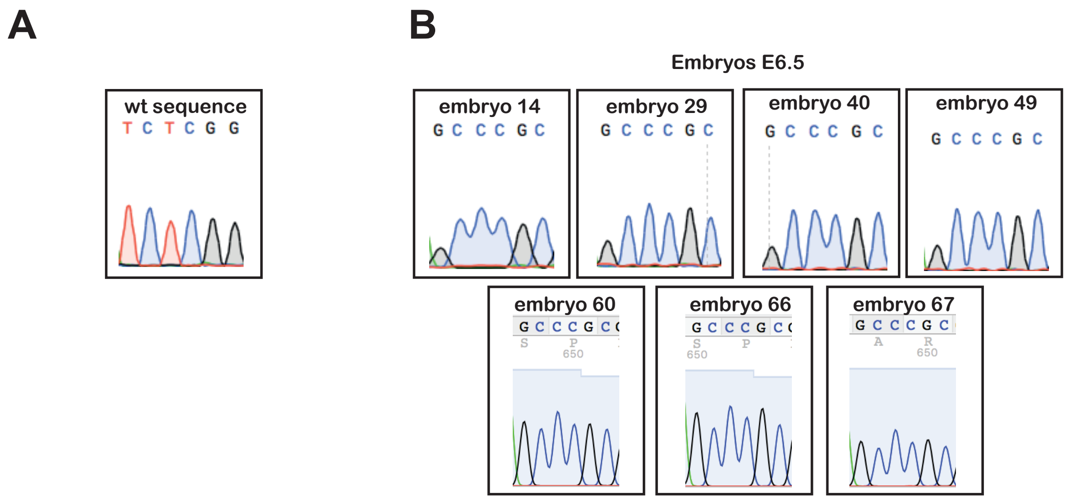


**Figure S3.** Sequences obtained from E6.5 embryos genotyped as homozygous for the S138A mutation by PCR. (A) Wild-type (wt) sequence. (B) Sequences from homozygous S138A mutant E6.5 embryos. The sequencing confirmed that all seven E6.5 embryos were homozygous.

**Figure S4**

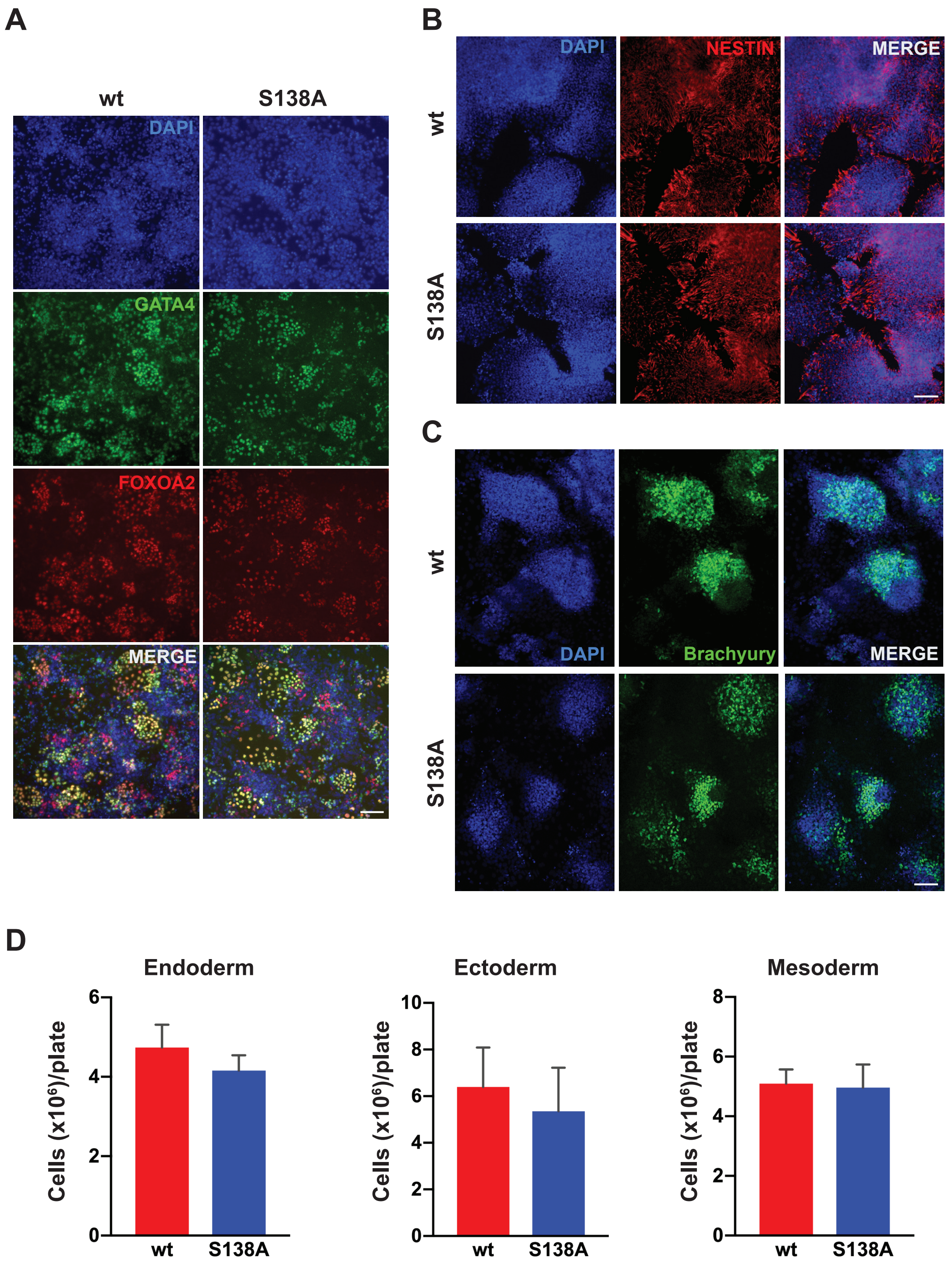


**Figure S4.** Differentiation to somatic lineages of wt and S138A ES cells. (A) Differentiation of ESC into endoderm. Wt and S138A cells were stained with anti-Gata4 and anti-FoxA2 after 3 days of differentiation. (B) Differentiation of ESC into ectoderm. Wt and S138A were stained with anti-Nestin antibody after 4 days of differentiation. (C) Differentiation of ESC into mesoderm. Wt and S138A were stained with anti-Brachyury antibody after 4 days of differentiation. All scale bars = 100 μm. (D) Quantification of the final number of cells per plate in the differentiation experiments (mean ± SEM, n=3).

**Figure S5**

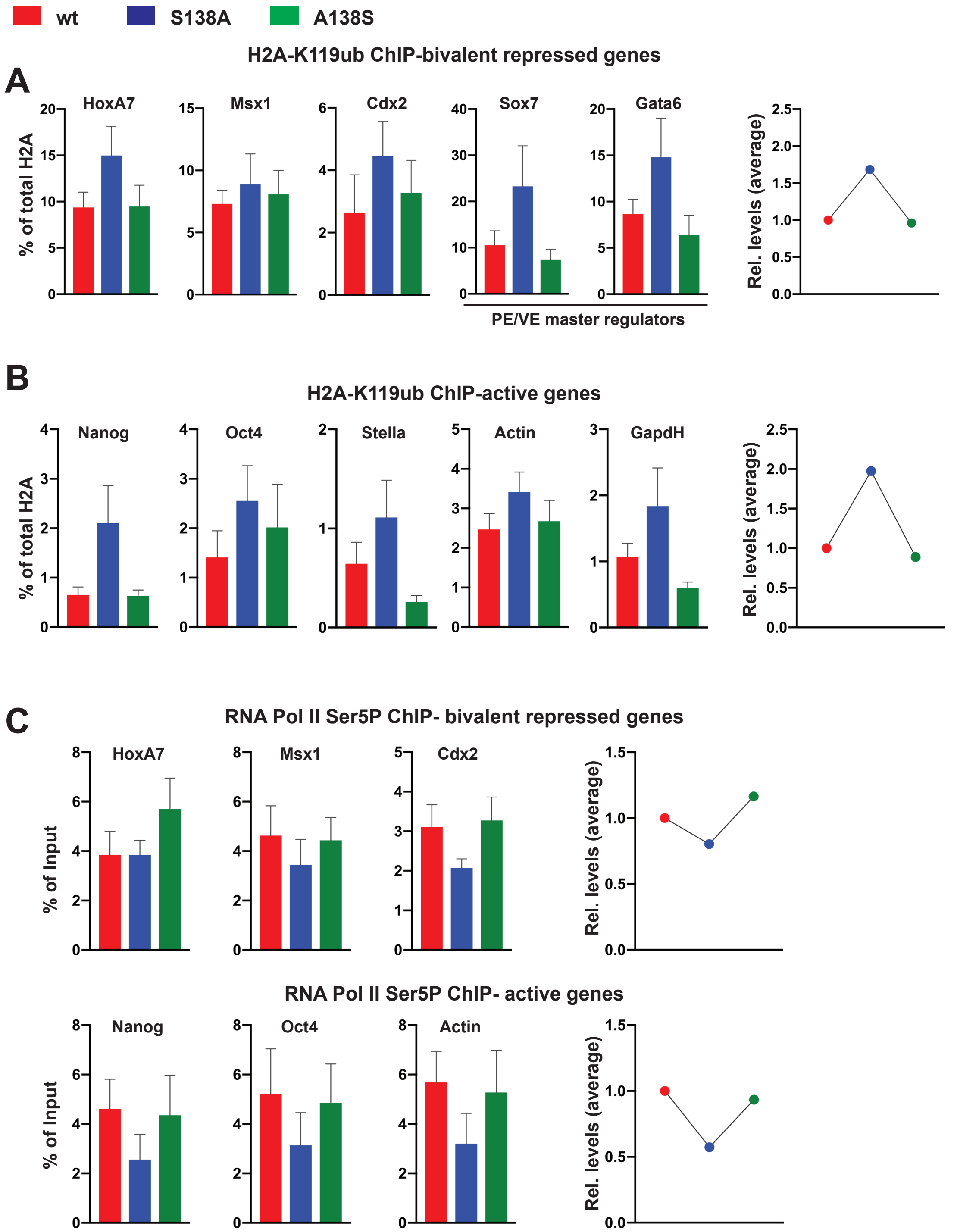


**Figure S5.** Changes in H2AK119Ub and Ser5Pol II levels in promoters of S138A mutant ESC. (A) ChIP-qPCR analysis of H2AK119Ub levels on bivalent promoters in wild-type (wt), S138A mutant and A138S revertant ESCs (n=6). (B) ChIP-qPCR analysis of H2AK119Ub levels in active genes promoters in wt, S138A and A138S ESCs (n=6). (C) ChIP-qPCR analysis of Ser5Pol II levels in bivalent promoters (top) and active genes (bottom) in wt, S138A and A138S ESCs. Graphs for individual promoters represent mean ± SEM, graphs on the right in each section represent the mean levels at the analysed promoters.

#### Table S1

Amino acid sequences of the C-terminal region of UBE2D3 from amniote and anamniote vertebrate species. The sequences shown cover residues 127-147. Ser138 in amniotes and Ala138 in anamniotes are shown in red and blue respectively. All other residues that differ from the human sequence are shown in green. The Acanthomorph sub-group of teleost fish are shaded in grey. Numbers next to the sequences indicate percentage identity with the human sequence for the C-terminal (C-term) and total sequences. Dashes indicate species where gaps in the

| Species | Common name | Amino acid sequence | C-term identity | Total identity |
|---|---|---|---|---|
| *Homo sapiens* | Human | K T D R D K Y N R I S R E W T Q K Y A M | | |
| *Pan troglodytes* | Common chimpanzee | K T D R D K Y N R I S R E W T Q K Y A M | 100% | 100% |
| *Gorilla gorilla* | Western gorilla | K T D R D K Y N R I S R E W T Q K Y A M | 100% | 100% |
| *Pongo pygmaeus abelii* | Orangutan | K T D R D K Y N R I S R E W T Q K Y A M | 100% | 100% |
| *Nomascus leucogenys* | Gibbon | K T D R D K Y N R I S R E W T Q K Y A M | 100% | 100% |
| *Macaca mulatta* | Rhesus macaque | K T D R D K Y N R I S R E W T Q K Y A M | 100% | 100% |
| *Macaca fasicularis* | Crab-eating macaque | K T D R D K Y N R I S R E W T Q K Y A M | 100% | 100% |
| *Papio anubis* | Baboon | K T D R D K Y N R I S R E W T Q K Y A M | 100% | 100% |
| *Chlorocebus sabeus* | Green monkey | K T D R D K Y N R I S R E W T Q K Y A M | 100% | 100% |
| *Callithrix jacchus* | Common marmoset | K T D R D K Y N R I S R E W T Q K Y A M | 100% | 100% |
| *Tarsius syrichta* | Tarsier | K T D R D K Y N R I S R E W T Q K Y A M | 100% | 100% |
| *Saimiri bolivensis* | Squirrel monkey | K T D R D K Y N R I S R E W T Q K Y A M | 100% | 100% |
| *Microcebus murinus* | Mouse lemur | K T D R D K Y N R I S R E W T Q K Y A M | 100% | 100% |
| *Otolemur garnetti* | Bushbaby | K T D R D K Y N R I S R E W T Q K Y A M | 100% | 100% |
| *Tupaia belangeri chinensis* | Chinese tree shrew | K T D R D K Y N R I S R E W T Q K Y A M | 100% | 100% |
| *Spermophilus tridecemlineatus* | Thirteen-lined ground squirrel | K T D R D K Y N R I S R E W T Q K Y A M | 100% | 100% |
| *Jaculus jaculus* | Lesser Egyptian jerboa | K T D R D K Y N R I S R E W T Q K Y A M | 100% | 100% |
| *Microtus ochrogaster* | Prairie vole | K T D R D K Y N R I S R E W T Q K Y A M | 100% | 100% |
| *Cricetulus griseus* | Chinese hamster | K T D R D K Y N R I S R E W T Q K Y A M | 100% | 100% |
| *Mesocricetus auratus* | Golden hamster | K T D R D K Y N R I S R E W T Q K Y A M | 100% | 100% |
| *Mus musculus* | House mouse | K T D R D K Y N R I S R E W T Q K Y A M | 100% | 100% |
| *Rattus norvegicus* | Norway rat | K T D R D K Y N R I S R E W T Q K Y A M | 100% | 100% |
| *Heterocephalus glaber* | Naked mole rat | K T D R D K Y N R I S R E W T Q K Y A M | 100% | 100% |
| *Cavia porcellus* | Gunea pig | K T D R D K Y N R I S R E W T Q K Y A M | 100% | 100% |
| *Chinchilla lanigera* | Chinchilla | K T D R D K Y N R I S R E W T Q K Y A M | 100% | 100% |
| *Octodon degus* | Brush tailed rat | K T D R D K Y N R I S R E W T Q K Y A M | 100% | 100% |
| *Oryctolagus cuniculus* | European rabbit | K T D R D K Y N R I S R E W T Q K Y A M | 100% | 100% |
| *Ochotona princeps* | Pika | K T D R D K Y N R I S R E W T Q K Y A M | 100% | 100% |
| *Sus scrofa* | Domestic pig | K T D R D K Y N R I S R E W T Q K Y A M | 100% | 100% |
| *Vicugna pacos* | Alpaca | K T D R D K Y N R I S R E W T Q K Y A M | 100% | 100% |
| *Manis pentadactyla* | Chinese pangolin | K T D R D K Y N R I S R E W T Q K Y A M | 100% | 100% |
| *Tursiops truncatus* | Common bottlenose dolphin | K T D R D K Y N R I S R E W T Q K Y A M | 100% | 100% |
| *Orcinus orca* | Killer whale | K T D R D K Y N R I S R E W T Q K Y A M | 100% | 100% |
| *Balaenoptera acutorostrata* | Minke whale | K T D R D K Y N R I S R E W T Q K Y A M | 100% | 100% |
| *Pantholops hodgsonii* | Tibetan antelope | K T D R D K Y N R I S R E W T Q K Y A M | 100% | 100% |
| *Bos taurus* | Cow | K T D R D K Y N R I S R E W T Q K Y A M | 100% | 100% |
| *Ovis aries* | Sheep | K T D R D K Y N R I S R E W T Q K Y A M | 100% | 100% |
| *Capra hircus* | Domestic goat | K T D R D K Y N R I S R E W T Q K Y A M | 100% | 100% |
| *Equus caballus* | Horse | K T D R D K Y N R I S R E W T Q K Y A M | 100% | 100% |
| *Ceratotherium simum* | White rhinoceros | K T D R D K Y N R I S R E W T Q K Y A M | 100% | 100% |
| *Felis catus* | Domestic cat | K T D R D K Y N R I S R E W T Q K Y A M | 100% | 100% |
| *Canis familiaris)* | Domestic dog | K T D R D K Y N R I S R E W T Q K Y A M | 100% | 100% |
| *Mustela putorius furo* | Ferret | K T D R D K Y N R I S R E W T Q K Y A M | 100% | 100% |
| *Odobenus rosmarus* | Pacific walrus | K T D R D K Y N R I S R E W T Q K Y A M | 100% | 100% |
| *Leptonychotes weddellii* | Weddell seal | K T D R D K Y N R I S R E W T Q K Y A M | 100% | 100% |
| *Pteropus alecto* | Black flying fox | K T D R D K Y N R I S R E W T Q K Y A M | 100% | 100% |
| *Pteropus vampyrus* | Megabat | K T D R D K Y N R I S R E W T Q K Y A M | 100% | 100% |
| *Myotis davidii* | David's myotis | K T D R D K Y N R I S R E W T Q K Y A M | 100% | — |
| *Myotis lucifugus* | Microbat | K T D R D K Y N R I S R E W T Q K Y A M | 100% | — |
| *Eptesicus fuscus* | Big brown bat | K T D R D K Y N R I S R E W T Q K Y A M | 100% | 100% |
| *Condylura cristata* | Star nosed mole | K T D R D K Y N R I S R E W T Q K Y A M | 100% | — |
| *Loxodonta africana* | African elephant | K T D R D K Y N R I S R E W T Q K Y A M | 100% | 100% |
| *Trichechus manatus* | Manatee | K T D R D K Y N R I S R E W T Q K Y A M | 100% | 100% |
| *Chrysochloris asiatica* | Cape golden mole | K T D R D K Y N R I S R E W T Q K Y A M | 100% | 100% |
| *Echinops telfairi* | Lesser hedgehog tenrec | K T D R D K Y N R I S R E W T Q K Y A M | 100% | 100% |
| *Orycteropus afer* | Aardvark | K T D R D K Y N R I S R E W T Q K Y A M | 100% | 100% |
| *Dasypus novemcinctus* | Armadillo | K T D R D K Y N R I S R E W T Q K Y A M | 100% | 100% |
| *Monodelphis domestica* | Opossum | K T D R D K Y N R I S R E W T Q K Y A M | 100% | 100% |
| *Sarcophilus harrisii* | Tasmanian devil | K T D R D K Y N R I S R E W T Q K Y A M | 100% | 100% |
| *Ornithorhynchus anatinus* | Platypus | K T D R D K Y N R I S R E W T Q K Y A M | 100% | — |
| *Falco cherrug* | Saker falcon | K T D R D K Y N R I S R E W T Q K Y A M | 100% | — |
| *Falco peregrinus* | Peregrine falcon | K T D R D K Y N R I S R E W T Q K Y A M | 100% | — |
| *Ficedula albicollis* | Collared flycatcher | K T D R D K Y N R I S R E W T Q K Y A M | 100% | — |
| *Zonotrichia albicollis* | White throated sparrow | K T D R D K Y N R I S R E W T Q K Y A M | 100% | 100% |
| *Geospiza fortis* | Medium ground finch | K T D R D K Y N R I S R E W T Q K Y A M | 100% | — |
| *Taeniopygia guttata* | Zebra finch | K T D R D K Y N R I S R E W T Q K Y A M | 100% | 100% |
| *Pseudopodoces humilis* | Tibetan ground jay | K T D R D K Y N R I S R E W T Q K Y A M | 100% | 100% |
| *Melopsittacus undulatus* | Budgerigar | K T D R D K Y N R I S R E W T Q K Y A M | 100% | 100% |
| *Anas platyrhynchos* | Mallard duck | K T D R D K Y N R I S R E W T Q K Y A M | 100% | — |
| *Gallus gallus* | Chicken | K T D R D K Y N R I S R E W T Q K Y A M | 100% | 100% |
| *Meleagris gallopavo* | Turkey | K T D R D K Y N R I S R E W T Q K Y A M | 100% | — |
| *Alligator mississippiensis* | American alligator | K T D R D K Y N R I S R E W T Q K Y A M | 100% | 100% |
| *Chelonia mydas* | Green sea turtle | K T D R D K Y N R I S R E W T Q K Y A M | 100% | — |
| *Chrysemys picta* | Painted turtle | K T D R D K Y N R I S R E W T Q K Y A M | 100% | 100% |
| *Pelodiscus sinensis* | Chinese softshell turtle | K T D R D K Y N R I S R E W T Q K Y A M | 100% | 100% |
| *Anolis carolinensis* | Carolina anole lizard | K T D R D K Y N R I S R E W T Q K Y A M | 100% | 100% |
| *Ophiophagus hannah* | King cobra | K T D R D K Y N R I S R E W T Q K Y A M | 100% | — |
| *Python bivittatus* | Burmese python | K T D R D K Y N R I S R E W T Q K Y A M | 100% | 100% |
| *Xenopus tropicalis* | Western clawed frog | K T D R E K Y N R I A R E W T Q K Y A M | 90% | 97% |
| *Xenopus laevis* | African clawed frog | K T D R E K Y N R I A R E W T Q K Y A M | 90% | 97% |
| *Nanorana parkeri* | Tibetan frog | K T D R E K Y N R I A R E W T Q K Y A M | 90% | 97% |
| *Ambystoma mexicanum-1* | Axolotl-1⌗ | K T D R E K Y N R I A R E W T Q K Y A M | 90% | — |
| *Ambystoma mexicanum-2* | Axolotl-2⌗ | K T D R A K Y N R I A R E W T Q K Y A M | 90% | |
| *Latimeria chalumnae* | Coelacanth | K T D R E K Y N R L A R D W T S K Y A M | 75% | — |
| *Tetraodon nigroviridis* | Tetraodon | K T D V T R Y T R T A K E W T A K Y A M | 60% | 91% |
| *Fugu rubripes* | Pufferfish | K T D V A R Y T K T A K E W T T K Y A M | 55% | 91% |
| *Oreochromis niloticus* | Nile tilapia | K T D S Q K Y T K M A K E W T Q K Y A M | 65% | 93% |
| *Neolamprologus brichardi* | Princess of Burundi | K T D P V R Y N K T A Q D W T Q K Y A M | 60% | — |
| *Astatotilapia burtoni* | Burton's mouthbrooder | K T D P V R Y N K T A Q D W T Q K Y A M | 60% | 91% |
| *Metriaclima zebra* | Zebra mbuna-a¶ | K T D S Q K Y T K M A K E W T Q K Y A M | 65% | 93% |
| *Metriaclima zebra* | Zebra mbuna-b¶ | K T D P V R Y N K T A Q D W T Q K Y A M | 60% | 91% |
| *Haplochromis nyererei* | Pundamilia nyererei-a¶ | K T D S Q K Y T K M A K E W T Q K Y A M | 65% | 93% |
| *Haplochromis nyererei* | Pundamilia nyererei-b¶ | K T D P V R Y N K T A Q D W T Q K Y A M | 60% | 91% |
| *Oryzias latipes* | Medaka-a¶ | K T D G Q K Y T K M A K E W T Q K Y A M | 65% | 93% |
| *Oryzias latipes* | Medaka-b¶ | K T D P V R Y N K T A Q D W T Q K Y A M | 60% | 91% |
| *Xiphophorus maculatus* | Southern platyfish-a¶ | K T D S Q K Y I K M A K E W T Q K Y A M | 65% | 93% |
| *Xiphophorus maculatus* | Southern platyfish-b¶ | K T D P V R Y N K T A Q D W T Q K Y A M | 60% | 91% |
| *Gasterosteus aculeatus* | Stickleback-a¶ | K T D N S R Y T K M A K D W T H K Y A M | 50% | 92% |
| *Gasterosteus aculeatus* | Stickleback-b¶ | K T D P V K Y N K T A Q D W T Q K Y A M | 65% | 92% |
| *Gadus morhua* | Atlantic cod | K T D L N K Y N K T A R E W T Q K Y A M | 75% | 95% |
| *Danio rerio* | Zebrafish | K T D T E K Y N R I A R E W T Q K Y A M | 85% | 96% |
| *Astyanax mexicanus* | Blind cave fish | K T D T E K Y N R I A R E W T Q K Y A M | 85% | 96% |
| *Lepisosteus oculatus* | Spotted gar | K T D R E K Y N R I A R E W T Q K Y A M | 90% | 97% |
| *Callorhinchus milii* | Elephant shark | K T D R E K Y N R I A R E W T Q K Y A M | 90% | 97% |
| *Petromyzon marinus* | Lamprey | K T D R E K Y N R I A R E W T Q K Y A M | 90% | 96% |
| *Branchiostoma floridae* | Lancelet (Chordata) | K T D K P R Y N E L A K E W T K K Y A M | 60% | 91% |
| *Ciona intestinalis* | Sea squirt (Chordata) | K N D R S K Y N D L A R E W T R K Y A T N | *65% | 86% |
| *Drosophila melanogaster* | Fruit fly (Arthropoda) | K T D R E K Y N E L A R E W T R K Y A M | 75% | 94% |
| *Culex tarsalis* | Mosquito (Arthropoda) | K T D R E K Y N E L A R E W T R K Y A M | 75% | 94% |
| *Corethrella appendiculata* | Midge (Arthropoda) | K T D R E K Y N E L A R E W T R K Y A M | 75% | 94% |
| *Zootermopsis nevadensis* | Dampwood termite (Arthropoda) | K T D R E K Y N E L A R E W T R K Y A M | 75% | 94% |
| *Lepeophtheirus salmonis* | Salmon Louse (Arthropoda) | K T D R S K Y M E M A R E W T K K Y A M | 70% | 88% |
| *Caenorhabditis elegans* | C. elegans (Nematoda) | K T D R E R Y N Q L A R E W T Q K Y A M | 75% | 93% |
| *Haliotis diversicolor* | Abalone sea snail (Mollusca) | K T D R Q K Y E D V A K E W T R K Y A M | 65% | 88% |
| *Fasciola hepatica* | Liver fluke (Platyhelminthes) | K T D R N K Y E D I A R E W T R K Y A M | 75% | 82% |
| *Saccharomyces cerevisiae* | Budding yeast (Ascomycota) | K T D R P K Y E A T A R E W T K K Y A V | 65% | 78% |
| *Schizosachaccharomyces pombe* | Fission yeast (Ascomycota) | K T D R S R Y E L S A R E W T R K Y A I | 60% | 84% |
| *Phanerochaete carnosa* | White-rot fungus (Basidomycota) | K T D R A R Y E A T A R E W T R K Y A M | 65% | 82% |
| *Hypsizygus marmoreus* | White beech mushroom (Basidomycota) | K T D R T R Y E A T A R E W T R K Y A M | 65% | 82% |
| *Acanthamoeba castellanii* | Acanthamoeba (Amoebozoa) | K T D R S R Y E N T A R E W T R K Y A M | 65% | 84% |
| *Arabidopsis thaliana* | Thale cress (Trachiophyta) | K T D K N K Y E S T A R S W T Q K Y A M G | *65% | 79% |
| *Galdieria sulphuraria* | Extremophilic red alga (Cyandiales) | K T N R G K Y E E T A R E W T R K Y A M | 60% | 85% |
| *Phyhophtera infestans* | Late blight water mould (Oomycota) | R T D R A R H D A T A R E W T A K Y A T | 50% | 81% |
| *Fistulifera solaris* | Oleaginous diatom (Bacillariophyta) | K T D R A R H D S T A R E W T S K Y A M | 60% | 81% |

**Notes on UBE2D3 sequences**

⌗Exon-6 and exon-7 were located for two Axolotl UBE2D genes by blast searching the Axolotl genome sequence (for details, see Supplementary Methods). Sequence comparison indicated that these genes were UBE2D2 and UBE2D3, but it was not possible to determine which is UBE2D3. Therefore, both sequences are shown as Axolotl-1 and Axolotl-2.

¶Duplicated UBE2D3 genes (UBE2D3-a and UBE2D3-b) arising from the whole genome duplication that occurred in the common ancestor to teleosts were detected in five species of Acanthomorph fish (Zebra mbuna, Pundamilia nererei, Medaka, Southern platyfish and Stickleback).

§The annotation of the UBE2D3 gene in the Carolina anole lizard was carried out by transcriptome mining (for details, see Supplementary Methods).

*Identities have been calculated for regions of homology and do not include the additional residue at the end of the sequence (Ciona intestinalis; Arabidopsis thaliana).

Values have been omitted for "Total Identity" where there were gaps in the sequence or where issues with the gene sequence meant that it was not possible to obtain a clear translated amino acid sequence (see Table S2).

**Table S2**

Gene co-ordinates/acccession numbers and sources for sequences shown in Figure 1 and Table S1

| Species | Common name | Gene co-ordinates or accession number | Source |
|---|---|---|---|
| *Homo sapiens* | Human | chr4:102797415-102826508 | UCSC genome browser |
| *Pan troglodytes* | Common chimpanzee | chr4:105343600-105372719 | UCSC genome browser |
| *Gorilla gorilla* | Western gorilla | chr4:114,153,960-114,185,280 | UCSC genome browser |
| *Pongo pygmaeus abelii* | Orangutan | chr4:107,249,729-107,279,260 | UCSC genome browser |
| *Nomascus leucogenys* | Gibbon | chr9:87,645,926-87,675,235 | UCSC genome browser |
| *Macaca mulatta* | Rhesus macaque | chr5:101,639,279-101,668,885 | UCSC genome browser |
| *Macaca fasicularis* | Crab-eating macaque | chr5:102,118,106-102,147,782 | UCSC genome browser |
| *Papio anubis* | Baboon | chr5:94,136,911-94,165,985 | UCSC genome browser |
| *Chlorocebus sabeus* | Green monkey | chr7:50,896,405-50,926,257 | UCSC genome browser |
| *Callithrix jacchus* | Common marmoset | chr3:90,826,499-90,855,726 | UCSC genome browser |
| *Tarsius syrichta* | Tarsier | KE935710v1:81,047-115,381 | UCSC genome browser |
| *Saimiri bolivensis* | Squirrel monkey | JH378138:12,313,524-12,343,189 | UCSC genome browser |
| *Microcebus murinus* | Mouse lemur | KQ054047v1:512,213-544,568 | UCSC genome browser |
| *Otolemur garnetti* | Bushbaby | GL873622:2,238,176-2,271,454 | UCSC genome browser |
| *Tupaia belangeri chinensis* | Chinese tree shrew | NW_006159808 | NCBI |
| *Ictidomys tridecemlineatus* | Thirteen-lined ground squirrel | NW_004936520.1 | NCBI |
| *Jaculus jaculus* | Lesser Egyptian jerboa | NW_004504368 | NCBI |
| *Microtus ochrogaster* | Prairie vole | NW_004949034.1 | NCBI |
| *Cricetulus griseus* | Chinese hamster | KE377574:1,794,741-1,819,159 | UCSC genome browser |
| *Mesocricetus auratus* | Golden hamster | NW_004801723 | NCBI |
| *Mus musculus* | House mouse | chr3:135439775-135465240 (MM10) | UCSC genome browser |
| *Rattus norvegicus* | Norway rat | chr2:240,627,638-240,654,620 | UCSC genome browser |
| *Heterocephalus glaber* | Naked mole rat | NW_004624830 | NCBI |
| *Cavia porcellus* | Gunea pig | scaffold_43:3,573,312,582,474 | UCSC genome browser |
| *Chinchilla lanigera* | Chinchilla | NW_004955496 | NCBI |
| *Octodon degus* | Brush tailed rat | NW_004524759.1 | NCBI |
| *Oryctolagus cuniculus* | European rabbit | chr15:46,797,666-46,830,252 | UCSC genome browser |
| *Ochotona princeps* | Pika | JH802128:6812054-6836211 | UCSC genome browser |
| *Sus scrofa* | Domestic pig | chr8:118,203,706-118,229,637 | UCSC genome browser |
| *Vicugna pacos* | Alpaca | KB632754:291,125-317,723 | UCSC genome browser |
| *Manis pentadactyla* | Chinese pangolin | KN013587:38,422-62,846 | UCSC genome browser |
| *Tursiops truncatus* | Common bottlenose dolphin | JH476740:35,239-60,565 | UCSC genome browser |
| *Orcinus orca* | Killer whale | NW_004438419 | NCBI |
| *Balaenoptera acutorostrata* | Minke whale | KI537457:2,977,101-3,003,691 | UCSC genome browser |
| *Pantholops hodgsonii* | Tibetan antelope | NW_005810708 | NCBI |
| *Bos taurus* | Cow | chr6:23358310-23369781 | UCSC genome browser |
| *Ovis aries* | Sheep | chr6:22121204-22146363 | UCSC genome browser |
| *Capra hircus* | Domestic goat | NC_030813 | NCBI |
| *Equus caballus* | Horse | chr3:37,154,007-37,167,562 | UCSC genome browser |
| *Ceratotherium simum* | White rhinoceros | JH767755:24,594,918-24,624,042 | UCSC genome browser |
| *Felis catus* | Domestic cat | chrB1:120,008,309-120,035,710 | UCSC genome browser |
| *Canis familiaris* | Domestic dog* | chr32:24,192,618-24,220,897 | UCSC genome browser |
| *Mustela putorius furo* | Ferret | GL896920:13,178,542-13,204,667 | UCSC genome browser |
| *Odobenus rosmarus* | Pacific walrus | NW_004450414 | NCBI |
| *Leptonychotes weddellii* | Weddell seal | NW_006383412 | NCBI |
| *Pteropus alecto* | Black flying fox | NW_006441953 | NCBI |
| *Pteropus vampyrus* | Megabat | scaffold_6805:5,409-33,086 | UCSC genome browser |
| *Myotis davidii* | David's myotis | NW_006297224 | NCBI |
| *Myotis lucifugus* | Microbat* | GL429896:2,318,363-2,333,718 | UCSC genome browser |
| *Eptesicus fuscus* | Big brown bat | NW_007370796 | NCBI |
| *Condylura cristata* | Star nosed mole | NW_004567122 | NCBI |
| *Loxodonta africana* | African elephant | scaffold_14:9,866,662-9,902,965 | UCSC genome browser |
| *Trichechus manatus* | Manatee | JH594677:3,950,779-3,988,061 | UCSC genome browser |
| *Chrysochloris asiatica* | Cape golden mole | NW_006408729 | NCBI |
| *Echinops telfairi* | Lesser hedgehog tenrec | JH980307:45,069,066-45,101,018 | UCSC genome browser |
| *Orycteropus afer* | Aardvark | NW_006921990 | NCBI |
| *Dasypus novemcinctus* | Armadillo | JH568248:2,922,267-2,951,557 | UCSC genome browser |
| *Monodelphis domestica* | Opossum | chr5:47129650-47175172 | UCSC genome browser |
| *Sarcophilus harrisii* | Tasmanian devil | chr6_GL864763_random:38475-79327 | UCSC genome browser |
| *Ornithorhynchus anatinus* | Platypus* | chrUn_DS181139v1:24317-26655 | UCSC genome browser |
| *Falco cherrug* | Saker falcon | NW_004994641 | NCBI |
| *Falco peregrinus* | Peregrine falcon | NW_004930191.1 | NCBI |
| *Ficedula albicollis* | Collared flycatcher | NC_021675 | NCBI |
| *Zonotrichia albicollis* | White throated sparrow | NW_005081558.1:7606696-7629775 | NCBI |
| *Geospiza fortis* | Medium ground finch | NW_005054304.1 | NCBI |
| *Taeniopygia guttata* | Zebra finch | NC_011467.1:21524003-21547354 | NCBI |
| *Pseudopodoces humilis* | Tibetan ground jay | NW_005087556.1:4655522-4675456 | NCBI |
| *Melopsittacus undulatus* | Budgerigar | NW_004848260.1:8888158-8911171 | NCBI |
| *Anas platyrhynchos* | Mallard duck | NW_004676847.1:548670-568729 | NCBI |
| *Gallus gallus* | Chicken | NT_455856.1 :8419869-8440682 | NCBI |
| *Meleagris gallopavo* | Turkey | chr4:40,307,474-40,314,153 | UCSC genome browser |
| *Alligator mississippiensis* | American alligator | NW_017709954.1 :369605-401191 | NCBI |
| *Chelonia mydas* | Green sea turtle | NW_006604451 | NCBI |
| *Chrysemys picta* | Painted turtle | NW_007281378.1: Region 107835-135019 | NCBI |
| *Pelodiscus sinensis* | Chinese softshell turtle | NW_005856091.1: Region 401358-428873 | NCBI |
| *Anolis carolinensis* | Carolina anole lizard | GAFD01029227.1 | NCBI |
| *Python bivittatus* | Burmese python | NW_006533296: Region 150799-174812 | NCBI |
| *Ophiophagus hannah* | King cobra | AZIM01000304 | GenBank |
| *Xenopus tropicalis* | Western clawed frog | chr1:64,170,894-64,188,573 | UCSC genome browser |
| *Xenopus laevis* | African clawed frog | chr1L:67,179,152-67,194,134 | UCSC genome browser |
| *Nanorana parkeri* | Tibetan frog | KN906220v1:1,038,792-1,080,495 | UCSC genome browser |
| *Ambystoma mexicanum* | Axolotl-1 | AMEXG_0030003997:702247-792645 | UCSC genome browser |
| *Ambystoma mexicanum* | Axolotl-2 | AMEXG_0030006217:1851620-1902148 | UCSC genome browser |
| *Latimeria chalumnae* | Coelacanth | JH130220:6637-59731+ | UCSC genome browser |
| *Tetraodon nigroviridis* | Tetraodon | chr10:12,247,820-12,251,495+ | UCSC genome browser |
| *Fugu rubripes* | Pufferfish | HE591832:182,178-186,047+ | UCSC genome browser |
| *Oreochromis niloticus* | Nile tilapia | chrLG19:9,932,884-9,937,895- | UCSC genome browser |
| *Neolamprologus brichardi* | Princess of Burundi | NW_006272052.1 | NCBI |

| | | | |
|---|---|---|---|
| *Astatotilapia burtoni* | Burton's mouthbrooder | NW_005179617.1 | NCBI |
| *Metriaclima zebra* | Zebra mbuna-a | >NC_036798.1:14907288-14914789 | NCBI |
| *Metriaclima zebra* | Zebra mbuna-b | NW_014444742.1 | NCBI |
| *Haplochromis nyererei* | Pundamilia nyererei-b | >NW_005187471.1:2672888-2679393 | NCBI |
| *Haplochromis nyererei* | Pundamilia nyererei-b | >NW_005187480.1:529850-536580 | NCBI |
| *Oryzias latipes* | Medaka-a | chr22:25,459,941-25,463,017 | UCSC genome browser |
| *Oryzias latipes* | Medaka-b | Chr10, >NC_019868.2:30876590-30885112 | NCBI |
| *Xiphophorus maculatus* | Southern platyfish-a | DNA scaffold: JH557024.1 | NCBI |
| *Xiphophorus maculatus* | Southern platyfish-b | >NC_036465.1:620591-625411 | NCBI |
| Gasterosteus aculeatus | Stickleback-a | chrXV:1410144-1416085 | UCSC genome browser |
| Gasterosteus aculeatus | Stickleback-b | chrIV:4895758-4906131 | UCSC genome browser |
| *Gadus morhua* | Atlantic cod | HE568491:62626-70609+ | UCSC genome browser |
| *Danio rerio* | Zebrafish | chr13:11614223-11630767 | UCSC genome browser |
| *Astyanax mexicanus* | Blind cave fish | KB871873.1:424117-432035 | ENSEMBL |
| *Lepisosteus oculatus* | Spotted gar | LG4:51219853:51242854:1 | ENSEMBL |
| *Callorhinchus milii* | Elephant shark | KI635875:786457-814714 | UCSC genome browser |
| *Petromyzon marinus* | Lamprey | GL477992:5467-35456 | UCSC genome browser |
| *Branchiostoma floridae* | Lancelet | C3ZGJ2 | Uniprot |
| *Ciona intestinalis* | Sea squirt | F6Y5B7 | Uniprot |
| *Drosophila melanogaster* | Fruit fly | P25867 | Uniprot |
| *Culex tarsalis* | Mosquito | A0A1Q3EUM9 | Uniprot |
| *Corethrella appendiculata* | Midge | U5EX56 | Uniprot |
| *Zootermopsis nevadensis* | Dampwood termite | A0A067R5A9 | Uniprot |
| *Lepeophtheirus salmonis* | Salmon Louse | C1BTS5 | Uniprot |
| *Caenorhabditis elegans* | C. elegans | P35129 | Uniprot |
| *Haliotis diversicolor* | Abalone sea snail | E6Y2Z0 | Uniprot |
| *Fasciola hepatica* | Liver fluke | A0A2H1C6N3 | Uniprot |
| *Saccharomyces cerevisiae* | Budding yeast | P15731 | Uniprot |
| *Schizosachaccharomyces pombe* | Fission yeast | P46595 | Uniprot |
| *Phanerochaete carnosa* | White-rot fungus | K5X3X9 | Uniprot |
| *Hypsizygus marmoreus* | White beech mushroom | A0A151VPJ7 | Uniprot |
| *Acanthamoeba castellani* | Acanthamoeba | L8GSE6 | Uniprot |
| *Arabidopsis thaliana* | Mouse ear cress | P35133 | Uniprot |
| *Galdieria sulphuraria* | Extremophilic red alga | M2XRN0 | Uniprot |
| *Phyoptera infestans* | Late blight water mould | Q8H728 | Uniprot |
| *Fistulifera solaris* | Oleaginous diatom | A0A1Z5J6L3 | Uniprot |

#### Table S3. Oligonucleotides used in this study.

| | Forward | Reverse |
|---|---|---|
| **Genotyping primers** | | |
| 1st PCR | TGCTGGCTGTTTGAAGAGGG | CCAGGCAGCTAACTCATTGGT |
| 2nd PCR wt specific | GTTTGTTTACAGGTACAACAGAATATCT | ATTCCAGCTATAATGCAGGTTATTC |
| 2nd PCR S138A specific | *ATCGTCTAA*GGTTTGTTTACAGGTACAACAGAATAGCC | |
| **ChIP-qPCR primers** | | |
| Msx1 | ACAGAAAGAAATAGCACAGACCATAAGA | TTCTACCAAGTTCCAGAGGGACTTT |
| HoxA7 | GAGAGGTGGGCAAAGAGTGG | CCGACAACCTCATACCTATTCCTG |
| Cdx2 | GGACTCCGCGAGCCAA | CTCAGCCCACGGTGCTC |
| Sox7 | TTTAGGGAAGTCAGTGCGCC | AGATCACACCCATGGCTTGG |
| Gata6 | GTTTCCCTCCCTCTTCTGCC | CGCTGTAACACATCCCCAGT |
| Nanog | CACAGTTTGCCTAGTTCTGAGG | GCAAGAATAGTTCTCGGGATGAA |
| Oct4 | GGCTCTCCAGAGGATGGCTGAG | TCGGATGCCCCATCGCA |
| Stella | AGAGCGGGGAATCCTACAGT | ACTGTAGGATTCCCCGCTCT |
| GapdH | CCACTTGTGGCAAGAGGCTA | GTGGAGAGTTGGGACGTGAG |
| β-Actin | GCAGGCCTAGTAACCGAGACA | AGTTTTGGCGATGGGTGCT |
| **Expression analysis primers** | | |
| Ube2d1 | GCAACCATCATGGGGCCCCC | CAAGACAAATACTCCCGTTG |
| Ube2d2 | TTCCACCATGGCTCTGAAGA | CTCCGCCCTGATAGGGACTA |
| Ube2d3 | ACAGACTATGGCGCTGAAAC | TCCCATAATTGTGGCTTGCC |
| MLN51 | ATGACGATGAGGATCGGAAAAAC | GTCCCCTTGGGTCGGACTTC |

**Table S4. Antibodies used in this study.**

| Antibody | Source | Application | Dilution |
|---|---|---|---|
| UBE2D3 | Abcam (Cat# ab176568) | WB | 1:3000 |
| UBE2D3-Ph | Frangini *et al.* 2013 | WB | 1:750 |
| H3 | Abcam (Cat# ab1791) | WB | 1:10000 |
| βTubulin | Cell Signaling (Cat# 2146) | WB | 1:2000 |
| HA | Cell Signaling (Cat# 3724) | WB | 1:1000 |
| CyclinD1 | Cell Signaling (Cat# 2922) | WB | 1:2000 |
| GFP | Cell Signaling (Cat# 2555) | WB | 1:2000 |
| H2A | Abcam (Cat# ab182255) | WB, ChIP | 1:2000, 2.5 μg |
| H2AUb | Cell Signaling (Cat# 8240) | WB, ChIP | 1:2000, 2.5 μg |
| Ser5Ph-RNA PolII | Biolegend (Cat# 904001) | ChIP | 2.5 μg |
| IgG | Diagenode (Cat# C15410206) | ChIP | 2.5 μg |
| Oct4 | Santa Cruz (Cat# sc-5279x) | IF | 1:200 |
| Gata6 | R&D Systems (Cat# AF1700) | IF | 1:100 |
| FoxoA2 | Abcam (Cat# ab108422) | IF | 1:400 |
| Gata4 | Santa Cruz (Cat# sc-1237) | IF | 1:300 |
| Nestin | Abcam (Cat# ab11306) | IF | 1:200 |
| Brachyury | Santa Cruz (Cat# sc-17743) | IF | 1:500 |
| PDGFRa-PE | eBioscience (Cat# 12_1401) | Flow Cyt. | 1:100 |
| PDGFR | Cell Signaling (Cat# 3174) | IF, PLA | 1:500 |
| FGFR1 | Cell Signaling (Cat# 9740) | IF, PLA | 1:200 |
| CBL | Novus (Cat# NBP 2-37574) | PLA | 1:200 |
| Tubulin-FITC | Sigma (Cat# F2168) | PLA | 1:50 |
| Alexa Fluor 488 Donkey anti goat | Invitrogen (Cat# A11055) | IF | 1:500 |
| Alexa Fluor 555 Donkey anti mouse | Invitrogen (Cat# A31570) | IF | 1:500 |
| Alexa Fluor 488 Donkey anti rabbit | Invitrogen (Cat# A32790) | IF | 1:500 |
| Alexa Fluor 594 Donkey anti rabbit | Invitrogen (Cat# R37119) | IF | 1:500 |
| Goat anti mouse-HRP | Invitrogen (Cat# A16084) | WB | 1:10000 |
| Goat anti rabbit-HRP | Invitrogen (Cat# A16096) | WB | 1:10000 |

**SUPPLEMENTARY METHODS:**

**Comparative analysis of UBE2D3 amino acid sequences**

Annotation of the UBE2D3 gene can present problems for automated gene annotation systems because of the high level of homology between different UBE2D family members. To address this problem, we made use of the fact that UBE2D3 in vertebrates has an alternative C-terminal exon (designated here as exon 7a, with amino acid sequence in amniotes: YNRLAREWTEKYAML) that is located between exon 6 and the predominantly expressed downstream exon 7b (sequence: YNRISREWTQKYAM). Exon-7a is not expressed in mouse ESC (data not shown). The fact that exon-7a is not present in other UBE2D family members and can be readily distinguished from the downstream C-terminal exon-7b by the presence of the additional lysine residue at the C-terminus allowed us to use exon-7a as a diagnostic feature to verify the identity of the UBE2D3 genes that we used for the vertebrate species comparison. All of the vertebrate UBE2D3 genes used to generate the sequences shown in Figure1 and Tables S1 and S2 were individually checked for the presence of exon-7a except for the Anolis carolinensis sequence, which was obtained by transcriptome mining (see below).

Quality assessment of the UBE2D3 sequences from fish

Due to the variable quality of the annotations of available fish genome sequences and the high level of sequence variation that was observed in the C-terminal exon in a subset of teleost species, all of the fish sequences shown in Tables S1 and S2 were checked manually by downloading the gene sequences, translating them and checking the sequences of the UBE2D3 exons. Where variant residues were found at intron/exon boundaries, the splice junctions were also checked manually to ensure that the variation was not caused by incorrect alignment of the splice sites. A number of the UBE2D3 sequences from amniotes and anamniotes also lacked the distally located amino terminal exon (MALKRINK in amniotes). Blast searching using available first exon sequences was used to search for the exon. In some species, it was not possible to locate the first exon (see Table S2). This is likely to be due to gaps in the available sequences. Files used for the sequence verification are available on request.

Annotation of the UBE2D3 sequence in the Carolina anole lizard (*Anolis carolinensis*).

We first used the Ensembl and UCSC gene annotation to check the status of position 138 of UBE2D3 in *Anolis carolinensis* and found that both databases annotated this position as alanine in the Anocar2.0 genome assembly. However, the ENSEMBL and UCSC databases rely on the first generation of gene annotation for the anole genome, which is primarily based on sequence homology with related species. This could lead to problems with the annotation due to the very high sequence conservation between UBE2D3 and UBE2D2. The fact that all of the other non-avian-reptile species and all avian species examined have serine at position 138 also suggested that the presence of the alanine might be due to a misannotation. We therefore carried out a further analysis using a new genome reannotation of *Anolis carolinensis* based on 14 adult and embryonic deep transcriptome (Eckalbar, et al. 2013). Our new analysis based on transcriptomic data in Eckalbar et al. shows that UBE2D3 with serine in position 138 is one of the genes that is poorly represented in the Anocar2.0 assembly because the contig with UBE2D3 cannot be mapped to Anocar2.0 with a high mapping rate. The second generation annotation for *A. carolinensis* ASU_Acar version 2.1 identified the UBE2D3 transcript as a8840. The sequence and ID of this transcript are currently available at NCBI (https://www.ncbi.nlm.nih.gov/nuccore/GAFD01029227.1). The transcript encodes a protein that is 100% identical to human UBE2D3 with serine at position 138. We conclude that the location predicted by the Ensembl and UCSC gene annotation as UBE2D3 is in fact UBE2D2 with alanine at position 138. We also conclude that UBE2D3 with serine at position 138 is expressed in *A. carolinensis* but was not annotated in Anocar2.0 because of incomplete genome assembly.

Identification of UBE2D3 in Axolotl (*Ambystoma mexicanum*)
The exceptionally large size of Urodele genomes (up to 10 times larger than humans) and the high levels of repetitive sequences in these genomes have made them difficult to sequence. An annotated genome sequence for Axolotl has recently become available (Nowoshilow, et al. 2018) and this allowed us to carry out a blast search for Axolotl UBE2D3 using the exon-6 and exon-7 sequences from the Tibetan frog (*Nanorana parkeri*). The blast search identified two UBE2D genes (for gene co-ordinates, see Supplemental Table 2). Using the co-ordinates, we were able to obtain the full sequence of exon-6 and exon-7 and the intron that separates them for both genes from the UCSC genome browser (https://genome.axolotl-omics.org). The intron was searched for the presence of the alternatively spliced exon-7a, but the alternative exon was not present in either gene sequence. The sequence of Exon-6 shows clearly that the genes are UBE2D2/3 and not UBE2D1/4. Examination of the intron sequence also shows that they are not duplicated as there is a ~3-fold difference in the length of the non-repetitive sequence in the introns of the two genes. We conclude from this that we have identified the UBE2D2 and UBE2D3 genes of Axolotl but the absence of the alternative Exon-7a does not allow us to say definitively which of the genes is UBE2D3. The sequence of Exon-7 differs between the two genes by one residue (A or E) at position 132. Therefore, the sequence of Axolotl UBE2D3 Exon-7 is shown as ambiguous for these residues in Figure 1 and Supplemental Tables 1 - 3. The absence of Exon-7a could be due to the exon having been lost during Axolotl evolution or it could be caused by a gap in the sequence.

#### Genotyping of point mutations in mouse ESCs and embryos
To genotype the point mutations, we used a nested allele-specific PCR (ASPCR), approach (Gaudet, et al. 2009). The mutated region was first amplified in a normal 15-cycle PCR. An aliquot of 1 µl from the first PCR was then subjected to a 25 cycle, 3-primer ASPCR with a common reverse primer and two genotype-specific forward primers. Both products can be discriminated due to a 10 base tail at the 5' end of the mutant specific primer. The final PCR products were run on a 3% MetaPhor™ agarose gel. The primers used for the genotyping are described in Table S5.

#### ChIP-qPCR analysis
ESCs were harvested, counted and diluted in medium to a final density of $10^6$ cells/ml prior to being crosslinked for 10 min at room temperature with 1%

formaldehyde. Crosslinking was quenched by adding glycine to 125 mM final concentration and incubating for 10 min at room temperature. After crosslinking, cells were washed 3 times with cold PBS. The resulting pellet was resuspended in 1.5 ml of swelling buffer (25 mM Hepes pH 7.5, 1.5 mM $MgCl_2$, 10 mM KCl, 0.1% NP40, Complete™ protease inhibitors cocktail), incubated in ice for 10 min and then homogenized in a dounce homogenizer (tight pestle). The nuclei were spun for 5 min at 3000g, 4°C and then resuspended in sonication buffer (50 mM Hepes pH7.5, 140 mM NaCl, 1 mM EDTA, 1% Triton-X100, 0.1% sodium deoxycholate, 0.1% SDS, Complete™ protease inhibitors cocktail) at a final density of $20x10^6$ cells/ml. Sonication was performed in a BioruptorPlus sonicator (Diagenode) for 30 to 45 min with 30 seconds on/off intervals to produce fragments of approximately 0.5-1 kb. Sonicated material was centrifuged for 10 min at 10000g, 4°C, to remove the insoluble fraction and supernantants were collected and snap frozen. Chromatin concentration and shearing of DNA were analysed by reverse crosslinking of an aliquot and running on an agarose gel.
Aliquots of 10 μg were used per immunoprecipitation (IP) and 1 μg used as input. For each IP, 2.5 μg of antibody (anti-H2A, anti H2Aub, anti Ser5RNAPolII or IgG) were incubated together with 15 μl protein A/G Dynabeads (ThermoFisher) in 1 ml of sonication buffer for 5 to 7 hours with gentle rotation at 4°C. Beads were washed once with sonication buffer and then the chromatin was added, IPs were performed O/N at 4°C with gentle rotation in a final volume of 1 ml. Antibody-bound chromatin was washed sequentially with 1 ml of the following buffers: sonication buffer, high salt buffer (50 mM Hepes pH 7.5, 500 mM NaCl, 1 mM EDTA, 1% Triton-X100, 0.1% sodium deoxycholate, 0.1% SDS), LiCl buffer (20 mM TriHCl pH8, 1mM EDTA, 250 mM LiCl, 0.5% NP40, 0.5% sodium deoxycholate) and TE buffer (10 mM TrisHCl pH8, 1 mM EDTA). After the washes, the chromatin was eluted from the beads and crosslinking was reversed by incubation in 200 μl of elution buffer (50 mM TrisHCl pH8, 50 mM NaCl, 1 mM EDTA, 1% SDS, 20 μg/ml DNAse-free RNAse A) for 6 hours at 68°C. Eluted DNA was treated O/N with 200 μg/ml proteinase K, at 42°C. Reverse crosslinking of the input was performed in parallel to that of the beads. DNA from input and IPs was purified using the QIAQuick PCR purification kit (Qiagen) following the manufacturer's instructions and eluted in 30 μl of water.
Real-time qPCRs were performed with Sensimix SYBR NORox SYBR GREEN (Bioline, UK) in a C1000 Thermal Cycler (BioRad). Each PCR was carried out in duplicate in a 20 μl final volume reaction with 400 nM primer concentration. Reactions contained 1 μl of the IPed DNA and 0.1 μl of the input DNA.
The oligonucelotides used for ChIP are listed in Table S5.

**RNA extraction and qPCR**

Total RNA was extracted from $1x10^6$ cells using Trizol (ThermoFisher) following the manufacturer's instructions. Reverse transcription was performed on 1 μg of RNA for each sample with the SuperScript II reverse transcriptase (ThermoFisher) according to the manufacturer's protocol. To quantify the levels of UBE2D family members mRNA, Real-time qPCRs were performed on the obtained cDNA using Sensimix SYBR NORox SYBR GREEN (Bioline). Each PCR was done in duplicate using 1 μl of a 1:10 dilution of cDNA and 400 nM primer concentration. The primers used in this analysis are listed in Table S5.

**ESC differentiation to germ layers**

*ESC differentiation to endoderm*: 10000 cells/cm$^2$ cells were plated on gelatin-coated plates and incubated in high glucose DMEM with 15% FBS, 100 units/ml penicillin, 100 μg/ml streptomycin, 0.1 mM non-essential amino acids, 1mM MTG, 1x GlutaMAX and supplemented with 25 ng/ml of FGF2 and 10 μM of retinoic acid for 3 days (Kim, et al. 2010).

*ESC differentiation to mesoderm*: cells were plated in gelatin + fibronectin-coated plates at a density of 15000 cells/cm$^2$ and incubated for 4 days in DMEM/F12 N2B27, 1x GlutaMAX, 100 units/ml penicillin, 100 μg/ml streptomycin, 0.1% β-ME and 30 ng/ml of Activin A.

*ESC differentiation to ectoderm*: cells were plated at a density of 15000 cells/cm$^2$ on gelatin-coated plates and incubated for 4 days in DMEM/F12-Neurobasal medium (50/50), 1x GlutaMAX, 100 units/ml penicillin, 100 μg/ml streptomycin, 100 μM βME, B27 minis Vitamin A and N2.

#### SUPPLEMENTARY DISCUSSION:

##### Effects of the whole genome duplication in teleosts

It is known that a whole-genome duplication took place in the common ancestor to all teleosts ~320-350 mya (Christoffels, et al. 2004; Vandepoele, et al. 2004). Genome duplications can have one of several different effects on the evolution of individual genes. One of the duplicated copies can be lost, both genes can remain the same, both genes can acquire sequence changes that divide the functions of the ancestral gene between them (subfunctionalization), or one gene can diverge and acquire a new function (neofunctionalization) (reviewed in (Glasauer and Neuhauss 2014). Duplicated UBE2D3 genes were initially identified in Medaka, Southern platyfish and Stickleback. The pattern of variation that was observed in the UBE2D3 genes that were initially identified in Cichlid fish did not follow the known phylogeny of Cichlids, suggesting that there might be additional unidentified duplicated UBE2D3 genes in these species. Blasting of Cichlid genomes with variant C-terminal sequences show that this was indeed the case. The blasting identified additional duplicated UBE2D3 genes in two Cichlid species (Zebra mbuna and Pundamilia nyererei). The duplicated genes are designated UBE2D3-a and –b (Figure 1, Table S1 and Table S2). These 5 species were the only Acanthomorph species where duplicated UBE2D3 genes could be confirmed, but it is likely that the presence of these genes is more widespread. More comprehensive sequencing will be required to confirm this. Comparison of the sequences reveals a significant divergence between of the duplicated genes in the C-terminal encoding region (Table S1) with the variable sequences located predominantly on the outward-facing side of the $\alpha$4-helix where they would be available for protein-protein contacts (Figure S1B). This suggests that sub-functionalization or neo-functionalization involving the generation of novel protein contacts are potential options for duplicated UBE2D3 genes.
